# Molecular computation at equilibrium via programmable entropy

**DOI:** 10.1101/2024.09.13.612990

**Authors:** Boya Wang, Cameron Chalk, David Doty, David Soloveichik

## Abstract

Synthetic molecular information processing is typically designed through programming kinetic pathways, so that molecules bind, unbind, or incur conformational changes in some desired order. In DNA nanotechnology, this paradigm is exemplified by strand displacement cascades that control signal transmission through toehold sequestering, and algorithmic tile assembly where correct computation emerges from enforcing ordered growth. In contrast, thermodynamic programming focuses solely on the desired end-state rather than the path, often allowing simpler reasoning and requiring fewer parameters, as well as avoiding energetically-favored, yet undesired, error states that often frustrate kinetic approaches. Here we demonstrate a thermodynamics-first paradigm based on the Thermodynamic Binding Networks (TBN) model, where the system’s equilibrium is determined by two simple factors: maximizing the number of complementary domain-level bonds while favoring configurations with more separate molecular complexes. We first validate the model by quantitatively measuring the free energy benefit of configurations with more separate complexes but identical binding, confirming the entropic driving force central to the TBN abstraction. We then construct signal propagation circuits including fan-in and fan-out, seeded-assembly systems that perform Boolean logic computation, and systems for synthesis of concatemers of size quadratic in that of the substrates (by computing their least common multiple). Realizing these TBN constructions required addressing experimental challenges including ensuring geometric feasibility of desired complexes through systematic domain ordering and compensating for multi-arm junction penalties. Our work may enable new ways to engineer complex molecular behaviors and help inform the understanding of the computational power of kinetics versus thermodynamics for molecular systems.

## 1 Introduction

Engineered molecular information processing has shown great promise with applications in nanoscience, synthetic biology, and therapeutics. As a biocompatible engineering material, DNA has been widely used for computation, diagnostics, nanorobotics, self-assembling structures, and super-resolution imaging tools^1–3^. Most of the molecular information processing systems are designed through programming a desired kinetic pathway: Molecules bind, unbind, or incur conformational changes to either release a kinetically-trapped output molecule, or assemble to the desired state through a specific binding order or partial order. Such kinetic programming matches the standard imperative programming practice of computer science. Although effective, such information processing often suffers from undesired side reactions that also occur, albeit at a slower rate. Mechanisms designed to reduce error usually rely on engineering error-correcting kinetic pathways, while errors are still thermodynamically preferred^4–7^. Engineering molecular systems that function correctly at thermodynamic equilibrium could, in contrast, have robust guarantees of correctness, since the desired state is also the energetically most favorable. Moreover and more broadly, programming algorithmic behavior encoded in thermodynamics could make programming certain tasks easier by alleviating the need to reason about exponentially-many possible kinetic pathways, or could enable entirely new types of computational behaviours. A key benefit enabled by thermodynamically-driven computing, compared to kinetically-driven systems requiring a specially prepared initial state, is that reusing the program with different inputs is straightforward no matter the underlying implementation: simply remove/deactivate the old input (e.g., add complement of input DNA strands), supply the new input, then wait for the system to reach its new equilibrium.

The first-principles understanding of the thermodynamics of Watson-Crick-Franklin base pairing has enabled modeling tools such as Mfold^8^, NUPACK^9^, oxDNA^10^, allowing the engineering of diverse thermodynamic functionalities. For example, molecular probes with carefully designed free-energies were adapted for molecular diagnostics such as differentiating single nucleotide polymorphisms^11^. Controlling the equilibrium of a DNA-based simple molecular mechanism allowed programming the phase transition of nanoparticles^12^. By engineering the activation energy of intermediate species, a reversible logic circuit was designed to respond to the change of inputs and recompute the output ^13^. Designing the equilibrium states of molecular systems at different temperatures allowed resetting molecular configurations through thermal cycling^14^. Carefully tuned weak interactions between pairs of strands have been shown to be capable of surprisingly complex digital and analog behavior^15^. However, we currently lack a sufficiently general abstraction at the thermodynamic level for systematic programming of systems with wide-ranging functionalities. Most designs are ad hoc for specific tasks, and understanding the thermodynamic behaviors relies heavily on the power of computational simulation, often at the computationally intensive level of individual nucleotides. To establish the design principles for systematically engineering molecular computation at thermodynamic equilibrium, it is necessary to develop a theoretical framework that is at the right level of abstraction such that the model is not far away from the underlying physics, but also simple enough for systematic investigation and rigorous mathematical proofs.

Recently the *thermodynamic binding networks* (TBNs)^16–23^ model has been proposed to aid in engineering functional molecules at thermodynamic equilibrium. In typical chemical systems, the thermodynamic equilibrium is determined by a large variety of chemical, structural, geometric, and combinatorial factors. The TBN model simplifies analysis of energy landscapes by coarsely modeling molecules in a well-mixed solution under a range of experimental regimes where certain model abstractions hold. First, bonds between complementary domains are strongly favored (and non-complementary domains are assumed to be *orthogonal*, i.e., to have zero binding affinity). Second, ties between multiple configurations with the same number of (domain-level) bonds are broken in favor of configurations with more separate complexes. In this work, the term ‘complex’ encompasses both multi-stranded structures and the degenerate case of a single-stranded DNA molecule. Intuitively, more separate complexes results in larger entropy due to additional microstates describing the position of each separate complex. Configurations which are maximally bonded and have the largest number of separate complexes among the maximally bonded configurations capture the thermodynamically preferred equilibrium in the model. This formalization of thermodynamic stability is valid in the regime in which bonds are too strong to dissociate spontaneously (e.g., DNA domains with 15 or more base pairs at room temperature), and also sufficiently dilute that the free energy of association penalty is significant. This simple energy abstraction makes it feasible to understand and analyze the robustness of molecular systems^24–26^. In addition, complex molecular behaviors have been theoretically constructed with thermodynamic guarantees of correctness, such as arbitrary Boolean circuit computation^16,18^, and engineering the intermediate configurations of the kinetic pathway in catalysis^19^.

The goal of this paper is to experimentally establish the TBN model as a design paradigm for engineering complex molecular systems. In Section 2.1, we start with the simplest non-trivial TBN system, demonstrating the feasibility of engineering the molecular equilibrium such that the maximally bonded configuration with the most separate complexes is strongly preferred. Then in Section 2.2, we engineer a signal propagation cascade that propagates information both “forward” and “backward”, including both fan-in and fan-out. Thermodynamic control also immediately accords new functionality to the same components: for example, fan-out in reverse generates AND logic. Then, in Section 2.3 we focus on algorithmic self-assembly, which, in contrast to non-algorithmic self-assembly such as DNA origami^27^, is typically programmed using imperative reasoning: Molecules are required to add in a particular order, or at least corresponding to the mathematical notion of a partial order^28^. In contrast we experimentally test a seeded algorithmic self-assembly system computing Boolean logic at thermodynamic equilibrium. This self-assembly design provides another way to compute any Boolean logic circuits as well as perform algorithmic self-assembly with provable guarantees of correctness^18^.

Finally, in Section 2.4 we develop a novel method to engineer concatemers (DNA strands consisting of repeated domains) with precisely defined size. We show that a “least common multiple” computation embedded in the thermodynamics of our system can be used to ligate a specific number of short concatemers together, allowing the total size of the resulting concatemer to be quadratic in the sizes of the starting concatemers. By programming short and cheap strands to assemble into long concatemers of specified size, our technique could be used, for example, as the template for the formation of periodic nano-structures.

In implementing the various TBN constructions we develop a methodology for faithfully realizing the TBN abstraction with DNA strands. In particular, we arranged domains on strands in an order which ensures geometric feasibility of resulting complexes (unpseudoknotted). Sequence design algorithms were used to ensure domain orthogonality and strong binding. Finally, when needed, we introduced single-base deletions to compensate for geometric energy penalties outside the scope of the TBN abstraction. By exploring the constraints the TBN model encountered for physical implementation, and finding methods to overcome these constraints, we provide a way to bridge the gap between idealized theory and realistic experimental implementation.

## 2 Results

### 2.1 The simplest TBN

The TBN model abstracts programming the thermodynamics of a chemical system in terms of the formal *enthalpy* and *entropy* of configurations, where enthalpy is equated with the number of bonds, and entropy with the number of separate complexes. Bonds are strictly favored—if one configuration has more bonds than another then the first is always preferred no matter how many more separate complexes the second has. A configuration that is maximally bonded is called *saturated*. The configuration with the largest number of separate complexes among saturated configurations is called *stable*. (TBN formal quantities of enthalpy and entropy do not unambiguously map onto physical enthalpy and entropy, but are intentional engineering simplifications. For instance, the free energy of base-pair formation has significant entropic terms, which would be lumped in TBN “enthalpy”. Further, note that no temperature dependence is considered.)

To use the TBN model as an effective programming paradigm, systems need to be engineered to satisfy the model’s abstraction. For DNA systems, TBN bonds correspond to domain-level Watson-Crick-Franklin base-paring and standard sequence design is performed to ensure orthogonal domain binding. Domain lengths are chosen long enough such that saturated configurations are strongly preferred. The order of domains has to be chosen to ensure feasible geometry of desired complexes (discussed later).

In this section, we study the arguably simplest non-trivial TBN containing four indivisible molecules *a, b, ab* and *a*^∗^*b*^∗^. (Abstract TBN molecules are drawn using the “box” notation in the figures. Note that prior TBN literature referred to indivisible molecules as *monomers* ^16,17^.) Bonds can form between starred and unstarred boxes with the same label (complementarity). The two configurations shown in Figure 1A have the maximum possible bonds and thus are saturated. The bottom configuration has three separate complexes compared with two for the top configuration. Since there are no saturated configurations with more separate complexes, the bottom configuration is stable according to the TBN model and is expected to be thermodynamically preferred.

**Figure 1:**
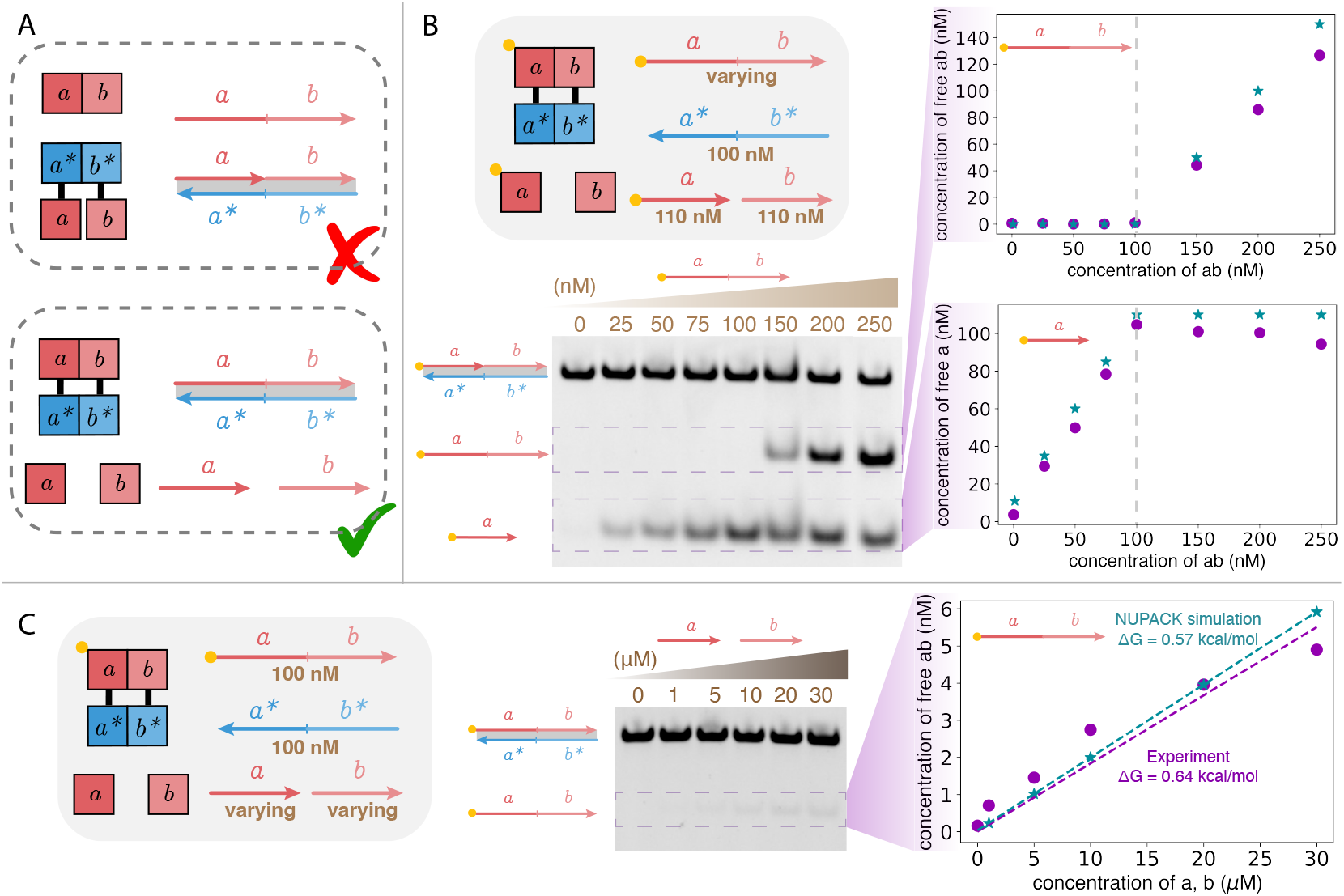
Experimental verification of the simplest non-trivial TBN. (A) Among the two saturated (maximum bond) configurations, the bottom configuration is stable (maximum bonds and also maximum number of separate complexes). Each box corresponds to a *domain* (consecutive DNA nucleotides) that can bind with its complementary domain (labeled by a star). (B-C) Unstained 12% native PAGE gel varying (B) strand *ab* or (C) strands *a* and *b* after annealing (95 °C to 25 °C within 1.5 h). The yellow dot represents the fluorophore. The concentration of each strand in the annealing reaction is labeled in the figure. The circled data are experimental measurements of band intensity in the gel after normalization (full gels with calibration controls are in Figure S1). The starred data are NUPACK simulation results (maximum complex size set to 3). In (B), a 10% excess of strands *a* and *b* was added to ensure that all starred domains are bound (strand *a*^∗^*b*^∗^) in case of pipetting or concentration-measuring errors. Likewise, to account for stoichiometric error in (C), complex *ab* : *a*^∗^*b*^∗^ was purified prior to mixing with *a* and *b* in order to ensure no excess of *ab*. The Δ*G* values shown are computed for 1 M concentration and temperature 25°*C*.

We call the free-energy difference of one separate complex “*one unit of entropy* “. For DNA strands, the standard free energy at 37 °C of associating two molecules is estimated as 1.96 kcal/mol^29^. At typical concentrations (250 nM, 37 °C) calculations show that one unit of entropy should be approximately equal to the enthalpy of a 8 base-pair bond^25^. The penalty of association can be further increased through reducing reaction concentration. Prior work has crucially used the entropic driving force maximizing the number of separate complexes to drive a variety of behaviors (e.g.,^30^). Thus it is expected that experimental systems and protocols can be designed to implement the TBN model with high fidelity.

To experimentally confirm that a single unit of entropy is sufficient to strongly differentiate the two configurations in Figure 1A at concentrations typical for DNA nanotechnology, we mapped the TBN to a set of DNA molecules as described in Section S1.1 and Section S4. As a proxy for thermodynamic equilibrium, we annealed the strands at concentrations shown in Figure 1(B)-(C) as described in the caption, and performed native polyacrylamide gel electrophoresis (PAGE). For greater sensitivity, we labeled strands *a* and *ab* by fluorophores; gel experiments without fluorophores were consistent with fluorophore-labeled gels (see Figure S1A for a comparison). Strands *a* and *ab* were labeled by identical fluorophores to ensure that any fluorophore-dependent energy contribution would be equal. As shown in Figure S1(B), we varied the amount of *ab*, confirming that up to 100 nM (the concentration of strand *a*^∗^*b*^∗^), there is almost no measurable free *ab*, while free *a* linearly increases up to saturation. These experimental results are close to the NUPACK^9^ prediction of the same system. (NUPACK is a software analysis tool that uses the thermodynamic nearest-neighbor energy model^29^ to make predictions such as the concentrations of various complexes, or the minimum free energy structure, of interacting DNA strands.) These data confirm the strong thermodynamic preference for the bottom configuration in Figure 1A.

To more quantitatively measure the thermodynamic driving force of one unit of entropy, in Figure 1C we increased the amount of strands *a* and *b* to the point where free *ab* could be detected. Freeing 5% of *ab* required 3000 times higher concentration of strands *a* and *b* than *ab*. The standard free energy of association derived from these data (0.64 kcal/mol) is close to the effective value fitting NUPACK simulation (0.57 kcal/mol) (methods in Section S2.3). Note that this deviates from the theoretical standard free energy of association at 25 °C (1.89 kcal/mol), which we computed from the value of 1.96 kcal/mol at 37 °C reported by literature^29^ (calculation in Section S2.2). The discrepancy is likely caused by weak spurious interactions of strands and the free energy contribution of the nick in the three-stranded complex *a* : *b* : *a*^∗^*b*^∗^. Note that the discrepancy only results in 11% difference in predicted concentration at 250 nM.

### 2.2 Signal propagation circuits

Intermolecular signaling circuits are common in biology, and their recapitulation with first-principles design has been a powerful test-case for dynamic DNA nanotechnology. As a prototypical example, strand displacement cascades modulate the accessibility of the region used for the initiation of displacement (toehold sequestering), and can achieve intermolecular signal propagation and computation on digital and analog signals^1^. Toehold sequestering is a form of kinetic control resulting in metastable complexes. Indeed, the desired output is typically at odds with the thermodynamic equilibrium of the system resulting in so called “leak” (see refs.^25,26^ for exceptions). In contrast, systems designed through thermodynamics cannot rely on metastable complexes and the toehold sequestering mechanism. Nonetheless, we can engineer complex signal propagation including combinatorial logic using the TBN abstraction.

Based on a previously proposed TBN AND gate^16,17^, we experimentally implemented a circuit reversibly converting between the pair of signals *A* (*a*_1_*a*_2_) and *B* (*b*_1_*b*_2_), and signal *C* (*c*_1_*c*_2_). As shown in Figure 2A, the system has 3 stable configurations that are expected to be favored at thermodynamic equilibrium. Since among these configurations *C* can be free only if *A* and *B* are sequestered, this system implements AND gate behavior (*A* + *B* → *C*) in the forward direction, and fanout (*C* → *A* + *B*) in the backward. Note that incorrectly releasing signal strands incurs a unit of entropy penalty (Figure S2). Note that there is no direct interaction between signal strands and the information processing is conceptually similar to allosteric action: input binding results in change of configuration which indirectly influences whether the output molecule is bound. Importantly, there is no forced sequence complementarity or identity between the signal strands, allowing for composition into longer cascades taking the output as the input to a downstream circuit^16^.

**Figure 2:**
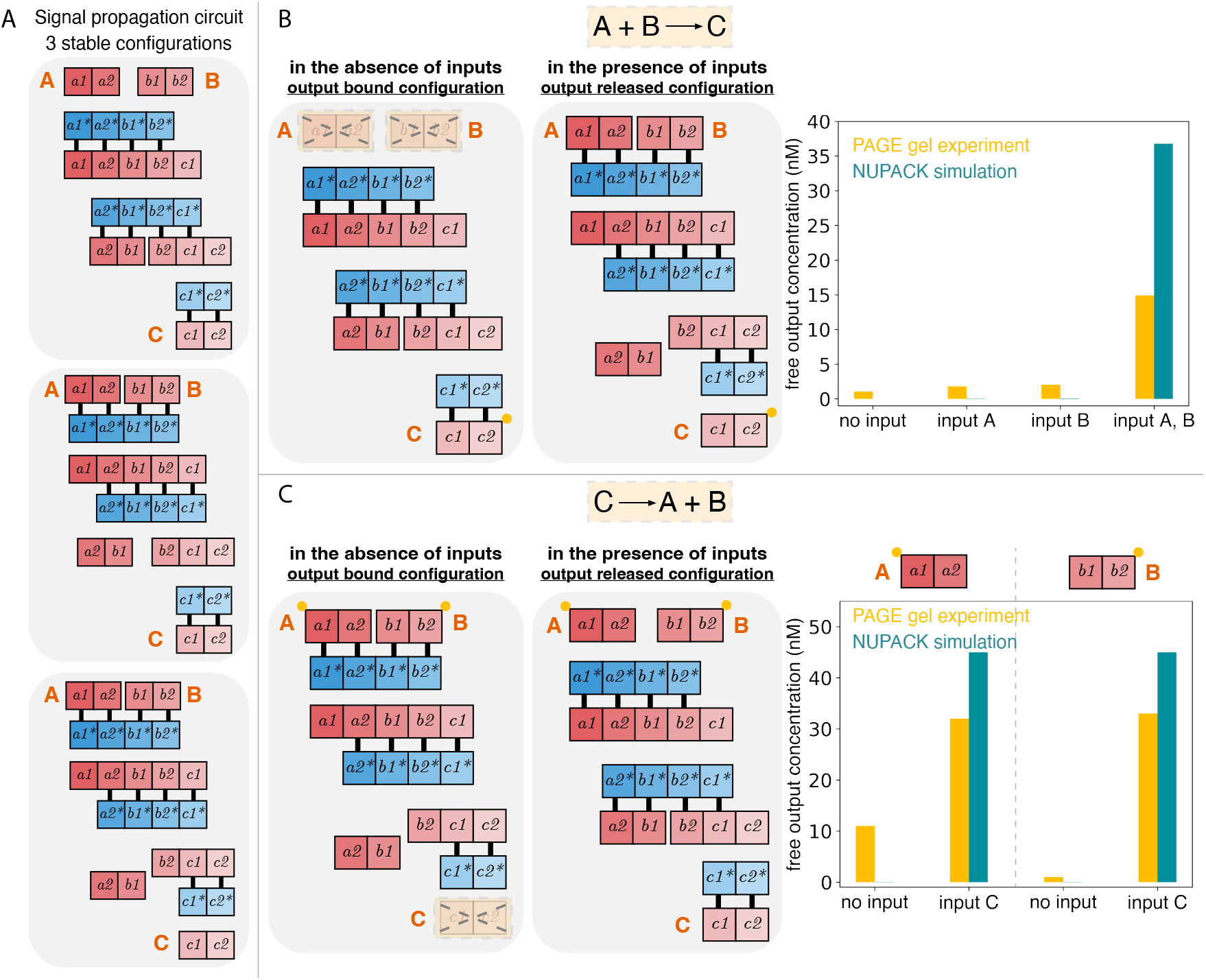
Experimental results for the signal propagation circuits. (A) The circuit contains 3 stable configurations. (B) Controlling the configuration of the circuits through adding the inputs *A* and *B*. The output *C* is labeled by a fluorophore (yellow dot). (C) Controlling the configuration of the circuits through adding the input *C*. In the experiment either output *A* or output *B* has a fluorophore label (yellow dot). All the species are 100 nM and slowly annealed from 90 °C to 25 °C with a rate of 0.1 °C per min. The yellow bar plots are experimental measurement of band intensity in native PAGE gels after normalization (full gels are in Section S3). The NU-PACK simulation (25 °C) used maximum complex size cutoff of 3.

To experimentally test the signal propagation circuit, we annealed all the strands with and without the inputs, and used native PAGE analysis to determine the amount of free output (fluorophore labeled). For the circuit implementing the reaction *A* + *B* → *C* (Figure 2B), the amount of free output *C* was 8 times more when both inputs *A* and *B* were present than when only one or none of the inputs was present. When we used the same circuit to implement the reaction *C* → *A* + *B* (Figure 2C), the amount of output was at least 3 times more when the input *C* was present. (In separate experiments either output *A* or output *B* had a fluorophore label.) The background signal could be due to imperfect stoichiometry between starred and unstarred strands caused by pipetting errors. In addition to gel experiments, we obtained similar results in fluorophore-quencher experiments in a plate reader (Figure S3B). NUPACK simulations confirm the overall pattern of the data, albeit with almost no background signal and somewhat higher correct free output concentration.

The middle stable configuration of Figure 2A has all signals sequestered; thus, depending on the concentrations of the constituent strands we expect less output signal(s) to be released than input signal(s) added. While we used identical concentrations of all strands, in principle the equilibrium concentration of free output can be increased by appropriately increasing the concentrations of non-input strands (Section S3.4). More sophisticated chemical information processing will require larger chemical circuits, and we explore potential scaling issues in the Discussion.

### 2.3 Seeded-assembling circuits

Both in evolved biology and engineered nanotechnology, molecular self-assembly is used to create complex structures from simple parts in a bottom-up manner^3^. For example, in DNA nanotechnology, DNA origami achieves the desired shape after annealing to thermodynamic equilibrium^31^. While the computational abilities of DNA origami are limited, algorithmic tile-assembly is effectively guided by algorithms encoded in molecular bonds^32,33^. Unlike DNA origami, algorithmic assembly is not thermodynamically favored over error pathways, and to prevent spurious binding between molecules precise control over binding kinetics—such as adjusting binding strength and temperature—is essential^32^.

As an example of *thermodynamic* algorithmic self-assembly, we experimentally realize self-assembling logic circuits proposed in prior work^18^. All the inputs are encoded by the choice of domains on a single (seed) strand, and the thermodynamic equilibrium determines the choice of other strands that bind and perform computation. With the input present, stable configurations of the TBN are guaranteed to have a self-assembled circuit with the correct logic (see Figure 3A).

**Figure 3:**
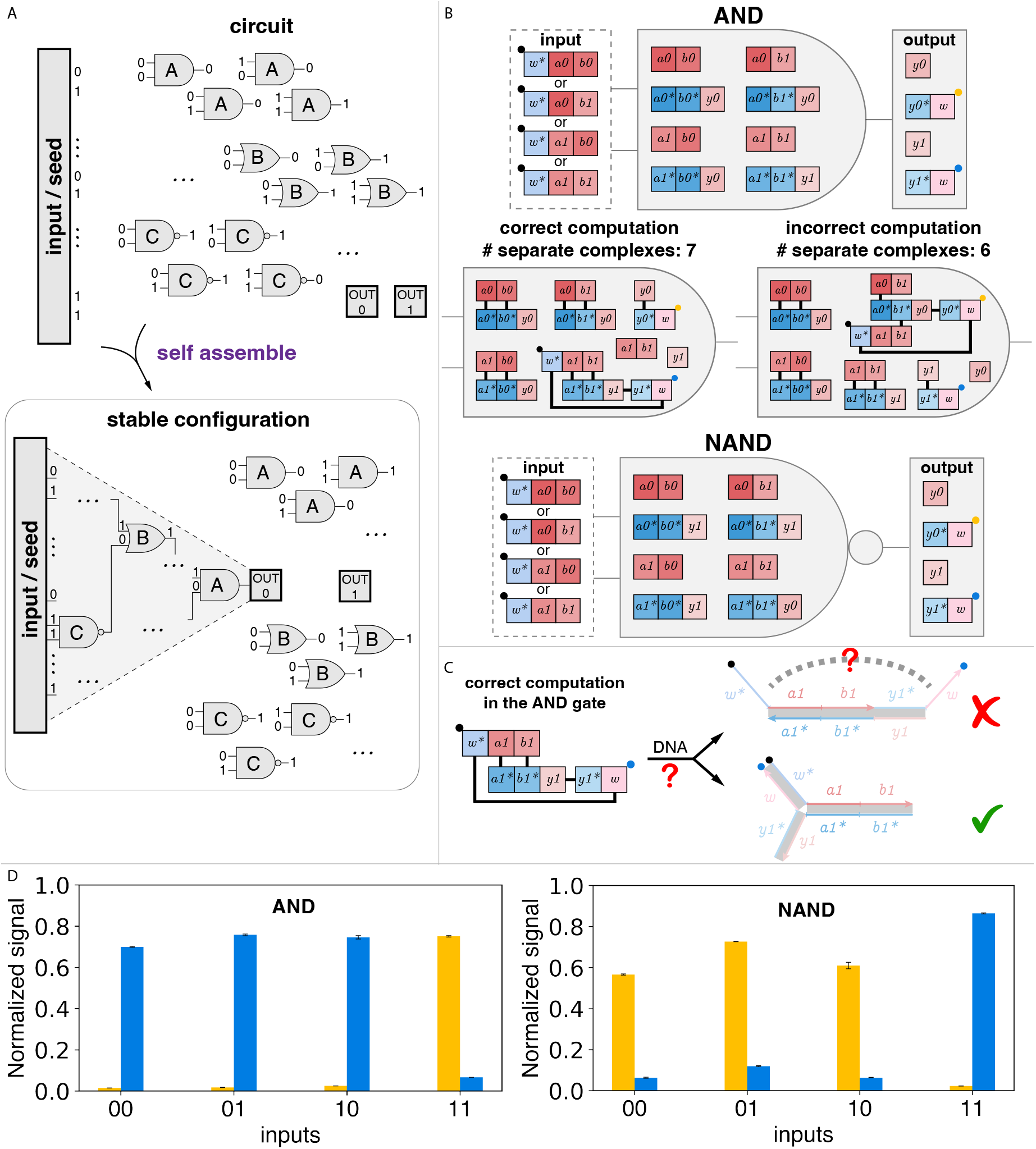
Experimental results for the seeded-assembling circuits. (A) Seeded-assembling circuits designed through the TBN model in principle have the thermodynamic guarantee that the stable configuration has the correct computation circuit. (B) Construction for the AND and NAND gates. The logic information for each gate is hard coded into the gate molecules. The input has a quencher attached, and each output strand has one of two different fluorophores. Which fluorophore is quenched is a proxy for which output is assembled with the input. As an example, the configuration of the correct computation for the input 11 has the output 1 in the same complex as the input, and incorrect computation has output 0 in the same complex as the input. The configuration with the correct computation has 1 more separate complex than any configuration with incorrect computation. (C) Converting the abstract TBN notation to DNA molecules may require organizing domain order in molecules to make DNA complexes geometrically feasible (methods in Section S4). (D) Fluorescence signals of the DNA implementation for a NAND gate and an AND gate with all inputs after annealing. Data were normalized according to the method in Section S5.3. It is not clear why why positive signals only reach 0.8. All species’ concentrations were 100 nM and slowly annealed from 90 °C to 25 °C with a rate of 0.1 °C per min.

We started with a 2-input 1-output AND gate and NAND gate (Figure 3B). The input strands have domains *w*^∗^, *a*_*i*_, and *b*_*j*_, with the two input values represented by *i, j* ∈ {0, 1}. The output strands have domains 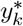 and *w*, with the output value represented by *k* ∈ {0, 1}. The logic value of the truth table of the gate is hard coded into the gate molecules 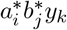. Strands *a*_*i*_*b*_*j*_ and *y*_*i*_, termed caps, induce a thermodynamic nucleation penalty which ensures that the only large complex that forms is with the seed (input). Each addition of a gate molecule onto a circuit which has its preceding matching gates present is entropy-neutral due to the displacement of the cap molecule from the gate molecule. The domain *w* creates a circular topology where completion of the circuit displaces an extra cap molecule, so that completion of the circuit is required in a stable configuration. The role of caps and the *w* domain is described further, with illustration, in Section S5.1. The output of the circuit is represented by which output molecule is in the same complex as the input. The correct computation always has one more separate complex than that of the incorrect computation (example of input 11 in Figure 3B). Correctness of both the logic and completion of the circuit can be proven formally^18^, and an abridged version of the proof is provided in section S5.2.

To measure the circuit output, we labeled the input strand by a quencher and the 0 and 1 outputs by two distinct fluorophores. Which of the two fluorescence signals is quenched indicates the result of the computation. To avoid potentially geometrically infeasible structures (Figure 3C), we followed a systematic procedure in determining the domain order on strands (Section S4). In each experiment, we annealed one chosen input with gates, caps and outputs and then measured the fluorescence signals for both fluorophores. We tested all possible inputs for both the AND and NAND gates. In each case the fluorescence signal for the correct output is almost fully quenched, and the fluorescence signal for the other output is greater than half of the maximum signal. (Maximum signal corresponds to the fluorescence of the output strands 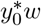 or 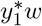 alone.)

We next scaled up self-assembling circuits, and in the process further explored TBN model non-idealities. One of the free energy contributions ignored by the TBN model is the penalty due to formation of multi-arm junctions. The previous one AND gate system only had a single 3-arm junction which we ignored in our design methodology. Can the TBN model still be used as an effective programming paradigm when more such junctions are unavoidable and their total free energy penalty could exceed one unit of entropy?

The self-assembling circuit construction with two cascaded AND gates (Figure 4) has two 3-arm junctions in the desired (assembled) configuration (Figure S13A). As a result, the free energy of the assembled structure is roughly 8-12 kcal/mol higher. To compensate, we introduced deletions in the “cap” molecules to increase the free energy of the configuration without assembly (Figure S13B). Deletions significantly improved the circuit performance compared to systems without deletions. Figure 4 shows the fluorescence output of the two AND gate construction for all eight inputs. For every input, the fluorescence signal for the correct output is about 20% (nearly fully quenched), and the fluorescence signal for the incorrect output is greater than 60% of the maximum signal. Note that a possible alternative approach, which we did not explore, involves adding a few “spacer” bases at the junctions to make them less geometrically constrained.

**Figure 4:**
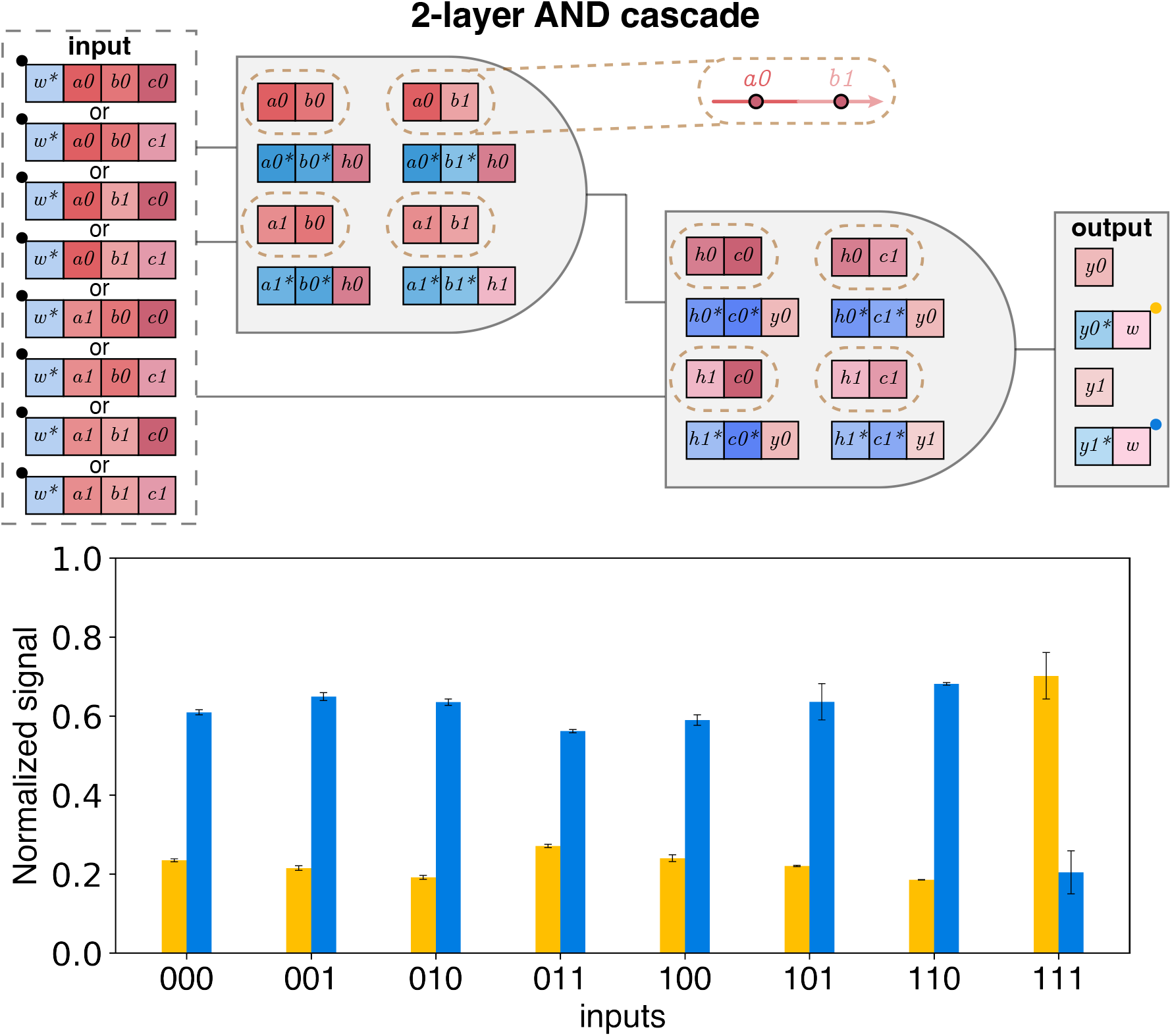
Construction and fluorescence signals for a seeded-assembling circuit of a 2-layer AND cascade with all inputs. The cap strands of the gates (highlighted by dashed lines) had single-base deletions; the deletions for *a*_0_*b*_1_ are in the call-out (red circles with black outline). Data were normalized according to the method in Section S5.3. All species were at 100 nM and slowly annealed from 90 °C to 25 °C with a rate of 0.1 °C per min.

### 2.4 Size-controlled concatemers

Synthetic nucleic acids have been used to construct finely controlled molecular structures^3^, and concatemers have been used as templates for engineering periodic structures^35^. Concatemers have been synthesized through enzymatic amplification^36^, or ligation of periodically hybridized DNA oligos, but it is hard to finely control the product size^37^. The apparent difficulty in self-assembling concatemers of programmed size can be articulated as follows: since the units are repeating, how does the system know when to stop growing?

Here we use the TBN model to construct concatemers of programmed size where the size is controlled by simple computation at thermodynamic equilibrium. The TBN construction contains two strands, composed of *m* and *n* repeated domains respectively. The domains between the two strands are complementary (*x* and *x*^∗^). When the ratio of the strands is *n* : *m*, the number of domains *x*^∗^ in the largest complex in the stable configuration is the least common multiple of *m* and *n*: LCM(*m, n*) (proof in Section S6). Note that the largest concatemer can be constructed from the smallest starting molecules when *m* and *n* differ by 1. In this way we can construct a concatemer of size quadratic in the sizes of starting substrates.

Figure 5A shows an example system with *m* = 3 and *n* = 2. To generate the concatemers, we annealed the two strands, followed by ligation and gel separation. To account for possible pipetting imprecision, we tested a range of concentration ratios by fixing the concentration of *x*^∗^*x*^∗^ and varying the concentration of *xxx*. Native PAGE gel confirmed that the products formed after annealing were sensitive to the initial concentrations (Figure 5B). After normalization, the phase change of the concentrations of the products as changing the initial *xxx* concentration is consistent with NUPACK prediction, and the peak of the desired complex is close to 200 nM—the ideal concentration ratio is 2 : 3. Note that we highlight two bands on the native gel to correspond to the desired product. Indeed, there are two different secondary structures with the same size satisfying the TBN model, one linear and one branched, which we expect are responsible for the two bands (Figure S15A). The denatured PAGE gel analysis shows the absence of products larger than the expected size. The smaller side product is hypothesized to be *x*^∗^*x*^∗^*x*^∗^*x*^∗^, caused by ligation of branched structure product (Figure S15).

**Figure 5:**
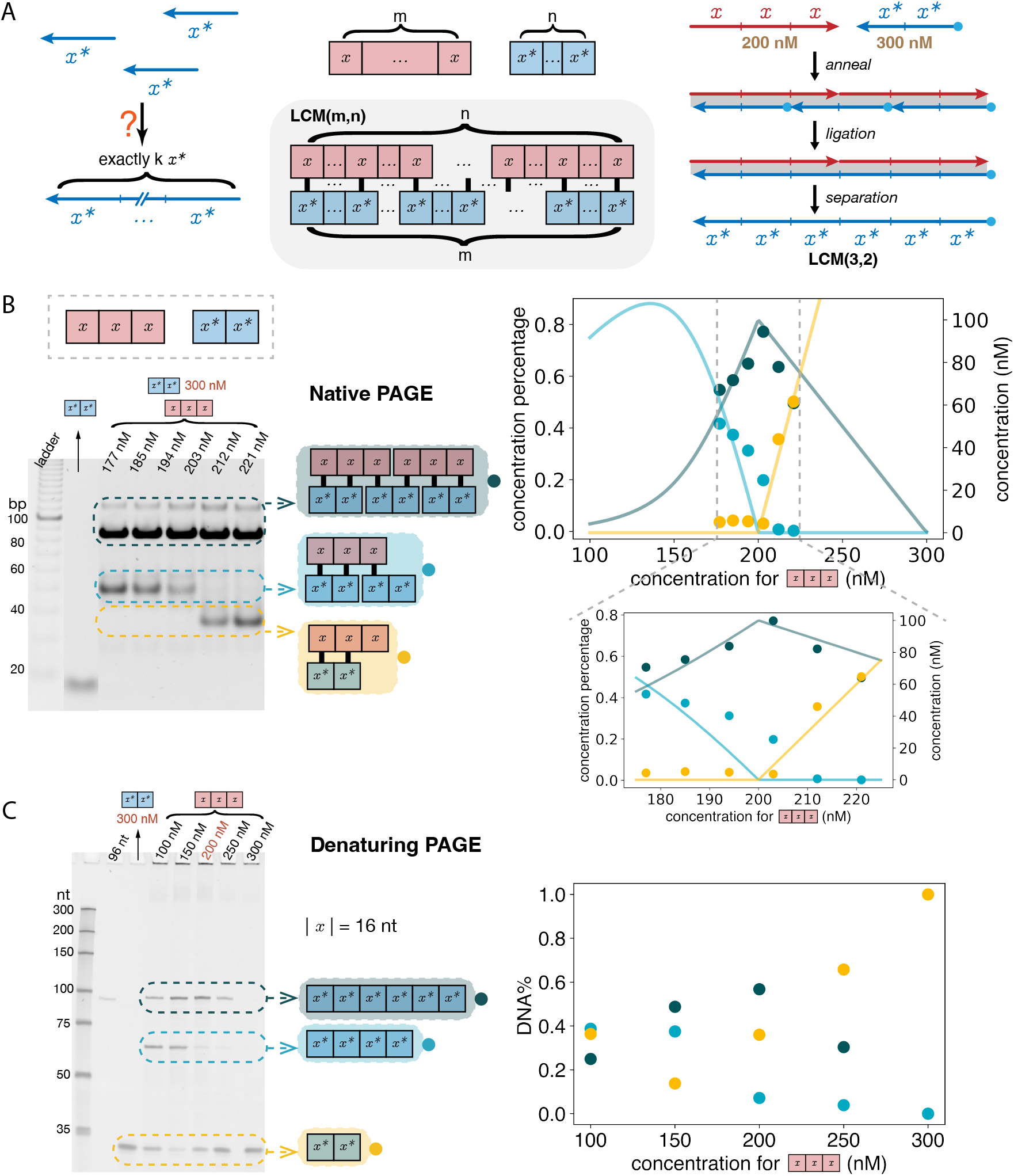
Size-controllable concatemers. (A) Unseeded self-assembly under the TBN model can be used to produce size-controllable concatemers. A TBN system with a pair of molecules with *m* and *n* repeated complementary domains has the stable configuration with the maximum possible number LCM(*m, n*) of domains for each molecule. As an example, when the ratio of *xxx* and *x*^∗^*x*^∗^ is 2 : 3, the stable configuration contains only the complex with LCM(3, 2) = 6 bound domain pairs. The 6-domain concatemer can be obtained after annealing, ligation and gel separation. The blue dot represents a phosphate label on 5’-end for ligation purpose. (B) The native 12% PAGE analysis of the annealed product with fixed *x*^∗^*x*^∗^ concentration and a range of different *xxx* concentrations. A range of concentrations of *xxx* were annealed with a fixed amount of *x*^∗^*x*^∗^ (300 nM). NUPACK simulation (line) is compared with the normalized native PAGE results (scatter plot). The color of the plots match the products showed in the gel. The NUPACK simulation has the maximum number of strands in a complex set to be 5. (C) The denaturing PAGE analysis of the ligated product after annealing a fixed amount of *x*^∗^*x*^∗^ with different concentrations of *xxx*. The *x* domain is 16 nt. Note that the *x* domain was composed of the ATC alphabet, which does not get stained by Sybr Gold ^34^, so it did not get imaged in the denaturing gel.

Next we generated different sizes of concatemers starting with substrates with different *m* and *n* values (Figure 6, Section S6.2). In every case, there was no apparent concatemer of length longer than the LCM. However, especially for longer target lengths, a substantial fraction of the total DNA was in shorter side-products. Further, when multiple pairs have the same LCM values (pairs (6, 3), (6, 2), and (3, 2)) starting with smaller substrates seemed to result in less DNA of full-length target size. Overall this TBN construction shows that a surprisingly simple computation performed by thermodynamic molecular self assembly can be effectively used to program concatemer assembly.

**Figure 6:**
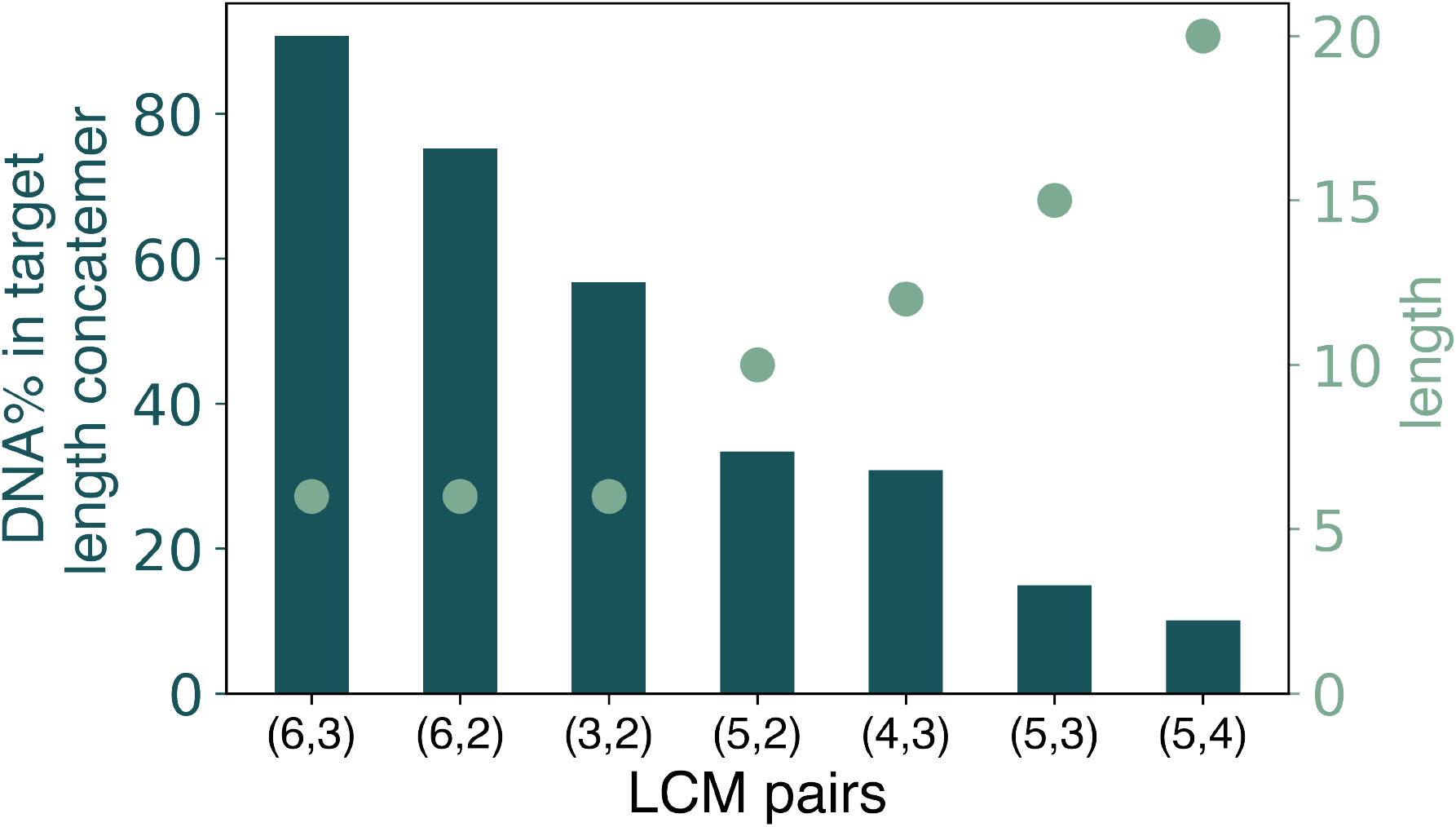
Generating size-controllable concatemers with different sizes of the substrate strands. Dots indicate the target size (LCM of the sizes of the substrates). The bars show the percent of DNA in the target size concatemer computed from band intensities in the denaturing gels (gels in Section S6.2).

## 3 Discussion

Much of experimental work in molecular programming focuses on kinetic approaches aiming to prescribe a desired sequence of molecular interactions (notable exceptions include DNA origami^31^ and non-complementary strand commutation networks^15^). However, kinetic approaches might unnecessarily shoehorn an ill-fitting programming paradigm motivated by the analogy to electronic computation where a program specifies a sequence of instructions. Unlike electronics, chemical systems naturally explore large configuration spaces with intrinsically random collisions of molecules. Here we have used an approach that embraces thermodynamic programming and leverages it as a fundamental design principle in the form of the TBN model. We designed and implemented a range of systems showing the flexibility of the approach: signal propagation circuits demonstrating conjunction and fan-out, seeded algorithmic self-assembly, and a system for assembling size-controlled concatemers based on the computation of least common multiple.

The demonstrated TBN constructions have thermodynamic guarantees relying on one unit of entropic difference (one separate complex) between desired and undesired behaviors. For example, in the signal propagation circuit, the output can be freed in the absence of the input at the cost of one entropy unit as shown in Figure S2. Similarly, the seeded-assembly circuit can assemble incorrectly also at the cost of one fewer separate complexes as shown in Figure 3(b). For larger systems—especially if the number of undesired configurations scales with the size of the system faster than the number of desired configurations— this one unit of entropy may not be large enough, resulting in a higher concentration of the undesired complexes. As an example, cascading signal propagation units increases the amount of leak (Section S3.4). Since one unit of entropy difference has a stronger effect at lower concentrations, one way to ensure desired behavior is via dilution. Nonetheless, since we must maintain reasonable experimental conditions, going beyond one unit of entropy is desired. Luckily, many previously described TBN designs, including for logic circuits and signal amplifiers, have a programmable units of entropy difference between the desired and undesired configurations^16,19,20^. It remains open, however, how other TBNs such as the self-assembly system can be made more robust in a similar way.

The TBN model applies to systems without geometric constraints, but using the model as an engineering paradigm requires considering geometric feasibility. We ordered domains on strands to ensure that desired structures are unpseudoknotted and introduced point deletions to compensate for the free energy penalty of multi-arm junctions. Alternatively, adding a couple of nucleotides at the base of the 3-arm junction could be an alternative way to stabilize it. More generally, adding “spacer” domains (e.g., poly-T regions) may be another approach to overcoming geometric constraints, and it may be feasible to allow pseudoknotted structures.

The original TBN model is specifically targeted at the regime where the enthalpic energy of forming a bond is strongly favored over the entropic energy of separating two molecules^16,17^, although more recent work allows for tuning the weight of enthalpic and entropic contributions in the model^19^. Recent experimental work has explored the computational behaviors achievable through designing weak bonds that bind different partners with various strengths^15^, yet the design principle of utilizing weak bonds to engineer systems with thermodynamic guarantees has not been thoroughly explored. In addition, thermodynamic equilibrium is a function of temperature. Incorporating consideration of temperature into the model could provide insights about engineering resettable DNA circuits through thermo-cycling. Programming the weight of both enthalpy and entropy and exploring the effect of temperature on preferred configurations would provide a better understanding of the thermodynamics in bioengineering and can lead to a more complete picture of programming thermodynamic systems.

A potentially important advantage of thermodynamics-first systems is their reusability. If the input is changed (e.g., if the original input strands are removed and new input strands are added), the thermodynamic equilibrium will naturally conform to the new input. Consider the contrast to systems whose correctness relies the interactions of meta-stable complexes: for example, DNA strand displacement systems are typically “use-once” and the initial DNA complexes have to be externally re-made in order for computation to proceed on a different set of inputs. Figure S12 shows input resetting in the NAND-gate self-assembly system. To switch the input, we deactivated the original input strand with its complement, and subsequently added the new input. (The significance of kinetic trace is described below.)

Throughout this work we have used an annealing protocol to drive the system to thermodynamic equilibrium in reasonable time. (It should be noted that certain systems do not readily converge to thermodynamic equilibrium even with slow annealing (e.g. HCR^38,39^) because the thermodynamically preferred configurations change during the process of annealing, allowing configurations to get “locked in” before becoming unfavorable. We believe that our systems anneal to equilibrium since the driving force separating the different configurations of interest is primarily entropic and thus temperature independent.) While thermal annealing is applicable for some applications such as paper-device sensing or molecular self-assembly, other applications require isothermal operation. For example, smart drugs are intended to operate within the body detecting and triggering on particular combinations of molecular cues^1^. Accordingly most of the strand displacement literature considers such a biologically compatible isothermal regime. We hypothesize that by partially destabilizing the domain binding we can achieve isothermal operation of TBNs. The kinetic trace in Figure S12 shows that the correct output can be produced on the order of minutes without annealing. Follow-up work is necessary to test the limits of input switching as well as to optimize the isothermal kinetics in a way that’s broadly applicable to TBNs.

Although a thermodynamics-first approach is established in many cases such as protein folding, it is not clear whether biological information processing can be more broadly understood as an equilibrium phenomenon. Nevertheless, de novo molecular engineering, though inspired by nature, does not have to follow nature’s “design principles” and might be more successfully inspired by areas where human engineering has particularly excelled.

A key lesson of decades of computer science is that clean abstractions—from Boolean circuits to high level programming languages—are necessary to create complex, reliable software systems. These abstractions restrict the design space (e.g., Boolean circuits restrict voltages to be high or low, and high-level programming languages prevent direct memory access) in a way that helps rather than hurts the design process. Counting the number of separate complexes provides a particularly clean abstraction of molecular systems, aligned with bottom up engineering practices. We have demonstrated through example systems doing signal propagation, Boolean logic, and synthesis of size-controlled concatemers, that this abstraction remains sufficiently expressive to achieve complex, yet reliable designed behavior.

## 4 Methods

Detailed experimental Materials and Methods are included in SI Appendix.

### 4.1 Annealing experiment

DNA oligonucleotides were purchased from Integrated DNA Technologies. DNA molecules were annealed to reach thermodynamic equilibrium. For the “simplest TBN,” the system was incubated at 95 °C for 5 minutes and then cooled at a rate of 0.1 °C/s to 20 °C. All other systems were incubated at 95 °C for 5 minutes and cooled at a rate 0.1 °C/min to 20 °C.

### 4.2 PAGE visualization

Novex 10% TBE precast gels (Invitrogen EC62752BOX) were used for the signal propagation circuit and the size controlled concatemer experiments. All other native PAGE gels were hand-cast with 1×TAE/Mg^2+^ buffer (0.04 M Tris, 1 mM EDTA, 12.5 mM Mg^2+^, pH balanced to 8.0 by acetate). The precast gels were run at 100 V for 10 mins and then 120 V for about 1 hour. The hand-cast gels were run at 180 V for about 3 hours at 25 °C. The denaturing gels were run at 100 V for 10 min and then 300 V for about an hour at 50 °C.

### 4.3 Fluorescence measurements

Fluorescence experiments were performed using the BioTech Synergy H1 multi-mode microplate reader. The excitation and emission wavelengths for the fluorophores were: 577 nm/608 nm for ROX, 555 nm/582 nm for ATTO550, 642 nm/668 nm for ATTO647. The excitation band-width was fixed at 9 nm and the emission bandwidth was fixed at 20 nm. Reads were taken every minute measured from bottom for at least half an hour to verify that equilibrium was reached and fluorescence signal would not change over time.

## Authors contributions

B.W. performed experiments and analyzed data. C.C. performed some experiments on the seeded-assembling circuit. D.D. proposed the size-controllable concatemer design. All authors discussed the results. B.W. wrote the initial draft of the manuscript. D.S. provided guidance throughout the project.

