## Supplementary Information for "Molecular computation at equilibrium via programmable entropy"

### Molecular computation at equilibrium via programmable entropy
### Supporting Information

Boya Wang[1,3], Cameron Chalk[1,4], David Doty[2], and David Soloveichik[1,*]

[1]*Electrical and Computer Engineering, University of Texas at Austin, Austin, TX 78712, USA*
[2]*Computer Science, University of California, Davis, CA 95616, USA*
[3]*Current address: Bioengineering, California Institute of Technology, Pasadena, CA 91125, USA*
[4]*Current address: Computation and Neural Systems, California Institute of Technology, Pasadena, CA 91125, USA*

#### Contents

**S1 Materials and Methods** **S3**
S1.1 Sequence design . . . . . S3
S1.2 DNA oligonucleotides . . . . . S4
S1.3 Annealing to reach thermodynamic equilibrium . . . . . S4
S1.4 Ligation experiment . . . . . S4
S1.5 PAGE visualization . . . . . S4
S1.6 Fluorescence measurements . . . . . S4
S1.7 NUPACK simulation . . . . . S5

**S2 The free energy of association from the simplest TBN system** **S6**
S2.1 Experiments verifying the stable configuration . . . . . S6
S2.2 Calculating the theoretical free energy of association at 25 °C . . . . . S7
S2.3 Deriving the free energy of association from data . . . . . S7
S2.4 Sequences . . . . . S8

**S3 Signal propagation circuits** **S9**
S3.1 Leak in the absence of inputs . . . . . S9
S3.2 Circuit for $A + B \to C$ . . . . . S10
S3.3 Circuit for $C \to A + B$ . . . . . S11
S3.4 Numerical analysis of equilibrium concentrations . . . . . S12
S3.5 Sequences . . . . . S15

**S4 Domain order for unpseudoknotted implementation** **S16**

**S5 Seeded-assembling circuits** **S19**
S5.1 The role of cap molecules and the $w$ domain . . . . . . . . . . S19
S5.2 Proof that stable configurations have correct circuits . . . . . . . . . . S20
S5.3 Data Normalization . . . . . . . . . . S22
S5.4 Kinetic pathway and input flipping . . . . . . . . . . S23
S5.5 Deletion . . . . . . . . . . S24
S5.6 Sequences . . . . . . . . . . S25

**S6 Size-controllable concatemers** **S28**
S6.1 Side product . . . . . . . . . . S28
S6.2 Concatemers with substrates of different sizes . . . . . . . . . . S29
S6.3 Sequences . . . . . . . . . . S32

### S1 Materials and Methods

#### S1.1 Sequence design

We first generated a pool of domains and then randomly mapped the sequences from the pool to the domains in the TBN systems. The domains were generated by satisfying the following:

1. To minimize self binding, (unstarred) domains are composed of only $A$, $T$, $C$.

2. To avoid synthesis errors, each domain does not contain 4 or more consecutive $C$'s, or more than 4 consecutive A's, or more than 4 consecutive T's.

3. To ensure the domains are approximately equally strong, the $C$ percentage in each domain is between $30\%$ to $42\%$.[1]

The concatenated strands also need to satisfy: no more than 4 consecutive A's or T's, no 4 or more consecutive G's, or no more than 7 consecutive A/T's.

We specified some design constraints with associated threshold values. Below, all references to "free energy" refer to NUPACK's definition of "complex free energy" (called "free energy of the complex" in [2]), the result of calling the function `pfunc` in the NUPACK Python interface. After repeatedly sampling the sequences and evaluating the scores based on the difference between calculated free energy and the threshold, the strands with better scores were selected. For sequence sets with similar scores, a sanity check was made to select the set with $> 99\%$ probability that strands form the desired structures in each system. The design constraints are the following:

1. **Domain-pair constraints**. Since the TBN model assumes that bonds form only between complementary domains, the domains in the pool need to be as orthogonal as possible. For a domain pair $\{a, b\}$, the free energy of 4 combinations of the domains and their complements ($\{a, b\}$, $\{a^*, b\}$, $\{a, b^*\}$, $\{a^*, b^*\}$) are compared with a threshold value $\alpha$.

2. **Single-strand constraints**. To reduce single strand's secondary structure, the free energy of each single strand is compared with a threshold $\beta$.

3. **Strand-pair constraints**. To reduce undesired binding between strands, the free energy of every strand pair is compared with a threshold $\gamma$. Note that $\gamma$ depends on the number of matching complementary domains in each strand, since strands containing complementary domains bind stronger.

$\alpha$, $\beta$ and $\gamma$ were chosen manually to optimize the tightness of the constraints and the run time.

Note that the sequences for generating concatemers has other special treatments: (1) when designing the sequences, the domain was cut into two orthogonal subdomains. Since the strands were composed of identical domains, if the domain contains non-negligible repeats, complementary domains could be misaligned when they bind. Cutting the domain into two orthogonal subdomains and designing sequences following the above method on the subdomain level could reduce the misalignment probability. (2) Every domain starts and ends with $C$ or $G$. Since it is possible to generate both a linear and a branched products (See Figure S15), the sequence at the nicks being $C$ or $G$ results in a more favorable stacking energy to favor the desired linear products.

The sequence design principle was implemented through the software Mathematica and *nuad* [3], and the code can be found at `https://github.com/boyawang-github/TBNexp`.

[1] To control domain binding strength, it is also common, and somewhat more accurate, to constraint the nearest-neighbor energy [1] of the domain binding to its complement, but we chose the simpler $C$ percentage heuristic.

#### S1.2 DNA oligonucleotides

DNA oligos were synthesized by Integrated DNA Technologies (IDT). Oligos were ordered purified in dry form and different purification options are listed in the corresponding sequence sections. We suspended the oligos in nuclease-free water (not DEPC-treated), and then quantified the concentration using $c = [\text{Absorbance}]/e$, where the 260 nm UV Absorbance value was measured through NanoDrop, and $e$ is the extinction coefficient provided by IDT. Usually, the nominal stock concentration was $\sim$100 $\mu$M.

#### S1.3 Annealing to reach thermodynamic equilibrium

The buffer for the concatemer experiment was $1\times$ T4 ligase buffer (diluted from $10\times$ T4 ligase buffer, NEB # B0202S). The buffer for all other experiments is TE/$Mg^{2+}$ buffer ($0.04$ M Tris, $1$ mM EDTA, $12.5$ mM $Mg^{2+}$, $0.01\%$ Tween 20, pH balanced to 8.0 by HCl). The annealing process was performed in a PCR thermocycler. For the simplest TBN system, DNA strands were incubated at 95 °C for 5 minutes and then slowly cooled down with rate 0.1 °C/s to 20 °C. For all other systems, DNA strands were incubated at 95 °C for 5 minutes and then slowly cooled down with rate 0.1 °C/min to 20 °C.

#### S1.4 Ligation experiment

After annealing, T7 ligase (NEB # M0318S) and fresh ATP (NEB # P0756S) were incubated with the annealed product at 25 °C for 30 min, adding sufficient volume of each so that the T7 ligase had final concentration 150 units/$\mu$L (from a stock of 3000 units/$\mu$L) and the ATP had final concentration 1 mM (from a stock of 10 mM). After ligation, the reaction products were inactivated by heat at 65 °C for 10 min. (Note that T4 ligase buffer is compatible with T7 ligase.)

#### S1.5 PAGE visualization

Novex 10% TBE precast gels (Invitrogen EC62752BOX) were used for the signal propagation circuit $C \to A + B$, and the size controllable concatemer experiments. All other native PAGE gels were homemade with $1\times$TAE/$Mg^{2+}$ buffer ($0.04$ M Tris, $1$ mM EDTA, $12.5$ mM $Mg^{2+}$, pH balanced to 8.0 by acetate). The running buffer was $1\times$TAE/$Mg^{2+}$. The precast gels were run using the XCell SureLock Mini-Cell Electrophoresis System at 100 V for 10 mins and then 120 V for about 1 hour. The homemade gels were run using the Hoefer SE600X Vertical Electrophoresis Systems at 180 V for about 3 hours at 25 °C.

Denature PAGE was run to separate and visualize the product of the concatemer. The denature PAGE gels were all homemade with $8$ M urea. The gels were run using the Hoefer SE600X Vertical Electrophoresis Systems at 100 V for 10 min and then 300 V for about an hour at 50 °C. The running buffer was $1\times$ TBE buffer.

We used Sybr Gold to stain the gels followed by scanning using Syngene G:Box imager. The excitation source was the UV illuminator and a standard Sybr Gold filter was used.

#### S1.6 Fluorescence measurements

Fluorescence experiments were measured on the BioTech Synergy H1 multi-mode microplate reader. The low volume NBS (non-binding surface) 384 well plates with clear flat bottom were used, purchased from Corning corporation (# 3544). The sample volume was chosen to be 20 $\mu$L, with a good signal-to-noise ratio, while minimizing total DNA (i.e., experiment cost). The excitation and emission wavelengths for the fluorophores are: 577 nm/608 nm for ROX, 555 nm/582 nm for ATTO550, 642 nm/668 nm for ATTO647. Throughout, the excitation bandwidth was fixed at 9 nm and the emission bandwidth was fixed at 20 nm. Reads were taken every minute measured from bottom for at least half an hour to verify that equilibrium was reached and fluorescence signal would not change over time.

##### S1.7 NUPACK simulation

NUPACK Python module [4] (version 4.0) was used for simulating experiments at equilibrium. The experiment temperature, sodium and magnesium concentration were set through the NUPACK command:
`Model(material=‘dna’, celsius=25, ensemble=‘stacking’,sodium=0.05, magnesium=0.0125)` The maximum number of strands in a complex is specified according to the configurations of each individual system.

### S2 The free energy of association from the simplest TBN system

#### S2.1 Experiments verifying the stable configuration

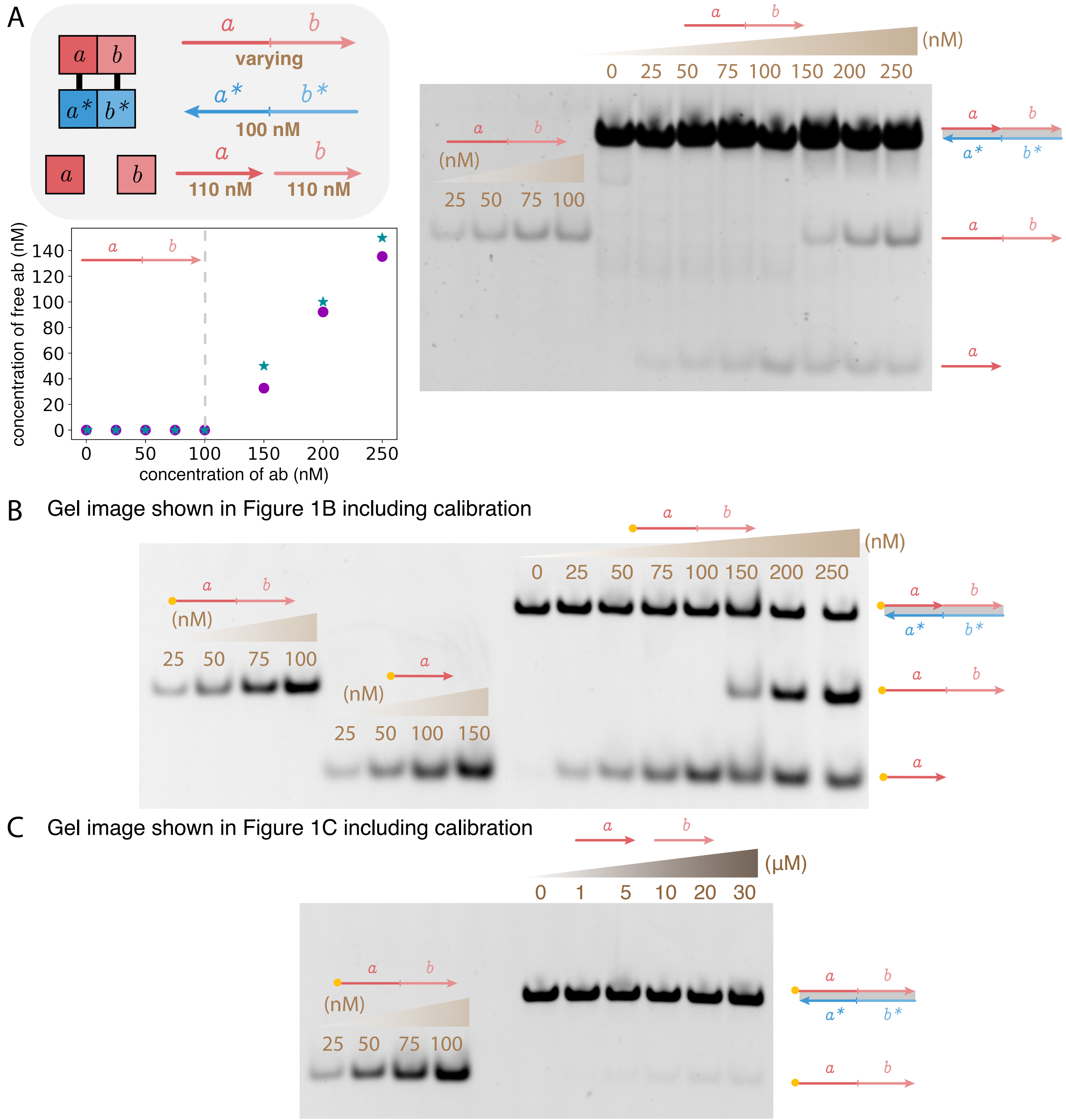


**Figure S1:** The same or similar experiments as Figure 1 investigating the simplest TBN system. (A) The experiment set up is the same as Figure 1B but without the fluorophore. The initial concentration of each molecule prior to annealing is labeled in the figure. The bands indicating the 3-stranded complex in the gel are of size within 40 to 50 bp according to a separate gel with a ladder (data not shown), which matches the desired size (45 bp) of the 3-stranded complex. The circled data are experimental measurement of band intensity in the gel after normalization. The starred data are NUPACK simulation. (B) and (C) show the corresponding 12% PAGE gel images in Figure 1B and Figure 1C including the calibrating species.

##### S2.2 Calculating the theoretical free energy of association at 25 °C

We calculate the standard (at 1 Molar) free energy of association at 25 °C ($\Delta G^\circ_{\text{assoc},25\ ^\circ\text{C}}$) according to the reported value (1.96 kcal/mol) at 37 °C ($\Delta G^\circ_{\text{assoc},37\ ^\circ\text{C}}$) [1]. The free energy of association should in principle be entirely entropic; however, the empirically reported value, which is used by NUPACK, has an enthalpic component of $\Delta H = 0.2$ kcal/mol. Treating the enthalpic component ($\Delta H$) and the entropic component ($\Delta S$) as independent of temperature, we use $\Delta G = \Delta H - T\Delta S$ to compute the free energy of association at 25 °C as $\Delta G^\circ_{\text{assoc},25\ ^\circ\text{C}} = \Delta H - \frac{T(25\ ^\circ\text{C})}{T(37\ ^\circ C)}(\Delta H - \Delta G^\circ_{\text{assoc},37\ ^\circ C}) = 1.89$ kcal/mol.

##### S2.3 Deriving the free energy of association from data

In this section we derive the free energy of association from the experimental data shown in Figure 1C. The system can be modeled as a chemical reaction with equilibrium constant $K$ as

$$a + b + ab : a^*b^* \overset{K}{\rightleftharpoons} a : b : a^*b^* + ab$$

Since the initial concentrations of $ab$ and $a^*b^*$ are equal (0.1 $\mu$M), we can view the molecule $ab : a^*b^*$ as having initial concentration $c = 0.1$ $\mu$M. Let the initial concentrations of $a$ and $b$ each be $x$. Calling the equilibrium concentrations of $a : b : a^*b^*$ and $ab$ as $y$, the equilibrium concentration of other species can be derived accordingly: $[a] = [b] = x - y$, and $[ab : a^*b^*] = c - y$. The concentrations satisfy

$$K = \frac{y^2}{(x-y)^2 \cdot (c-y)}.$$

Since we know the amount of free $ab$ at equilibrium is very small, we have $x \gg y$ and $c \gg y$. Thus $x - y \approx x$, $c - y \approx c$, and the above expression can be simplified as:

$$K = \frac{y^2}{x^2 \cdot c}$$

Simplifying for $y$, we can get:

$$y = \sqrt{cK} \cdot x$$

Thus when the amount of $y$ is small, $y$ linearly increases with $x$ (since $c$ and $K$ are constant).

The standard free energy can be derived from the slope of a linear fitting of $x$ (the initial concentration of $a$ and $b$) and $y$ (the equilibrium concentration of free $ab$): since the slope is $m = \sqrt{cK}$, we have $K = m^2/c$, and use the relationship $\Delta G^\circ = -RT \ln K$. As shown in Figure 1C, the standard free energy of association can be derived from both the experiment measurements and NUPACK simulation. The NUPACK simulation was conducted according to Section S1.7 using the same sequences as experiment, with the maximum number of strands in a complex being 3.

##### S2.4 Sequences

The unmodified oligos were ordered PAGE purified. The fluorophore labeled oligos were ordered HPLC purified.

| | |
|---|---|
| a | ATCTACTTATCAATCTATCTCTCTT |
| b | ACACAACACAAACCACAACA |
| ab | ATCTACTTATCAATCTATCTCTCTT ACACAACACAAACCACAACA |
| a*b* | TGTTGTGGTTTGTGTTGTGT AAGAGAGATAGATTGATAAGTAGAT |
| a-F | /56-ROXN/ATCTACTTATCAATCTATCTCTCTT |
| ab-F | /56-ROXN/ATCTACTTATCAATCTATCTCTCTT ACACAACACAAACCACAACA |

#### S3 Signal propagation circuits

##### S3.1 Leak in the absence of inputs

Figure S2 shows that generating free output $C$ in the absence of inputs $A$ and $B$ (i.e., leak) results in one fewer units of entropy (i.e., one fewer separate complex). Even with one input present and the other absent, the same unit of entropy penalty cost exists. This can be confirmed by TBN analysis tools such as stablegen [5, 6] or StableTBN [7, 8].

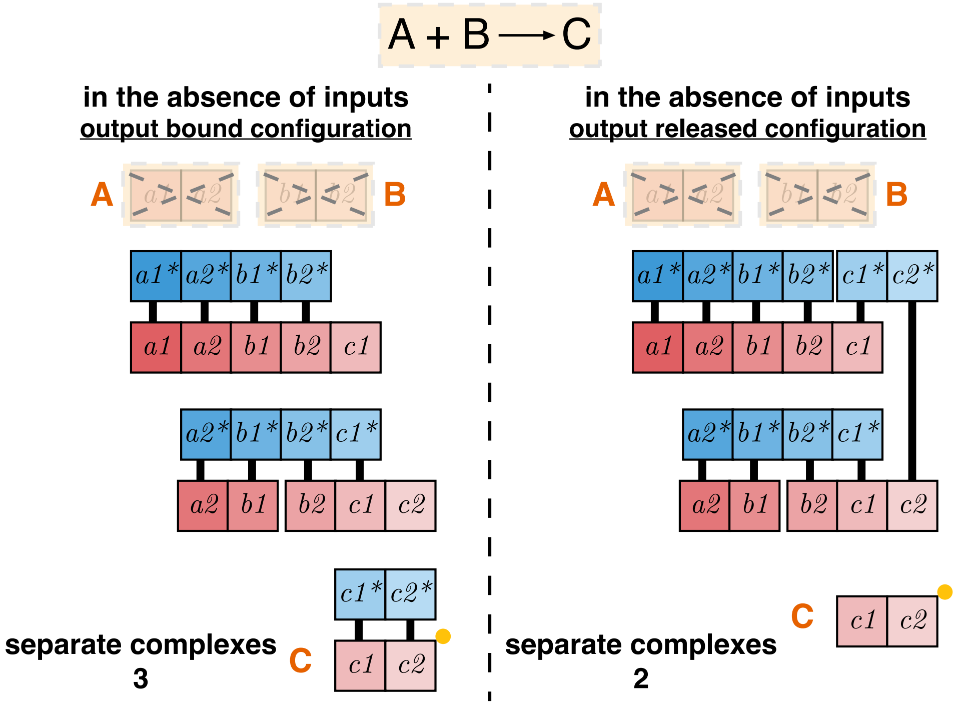


**Figure S2:** The configurations with output bound and unbound for signal propagation circuit $A + B \to C$ in the absence of inputs. To have the output released in the absence of inputs, there is one unit of entropy cost.

#### S3.2 Circuit for $A + B \to C$

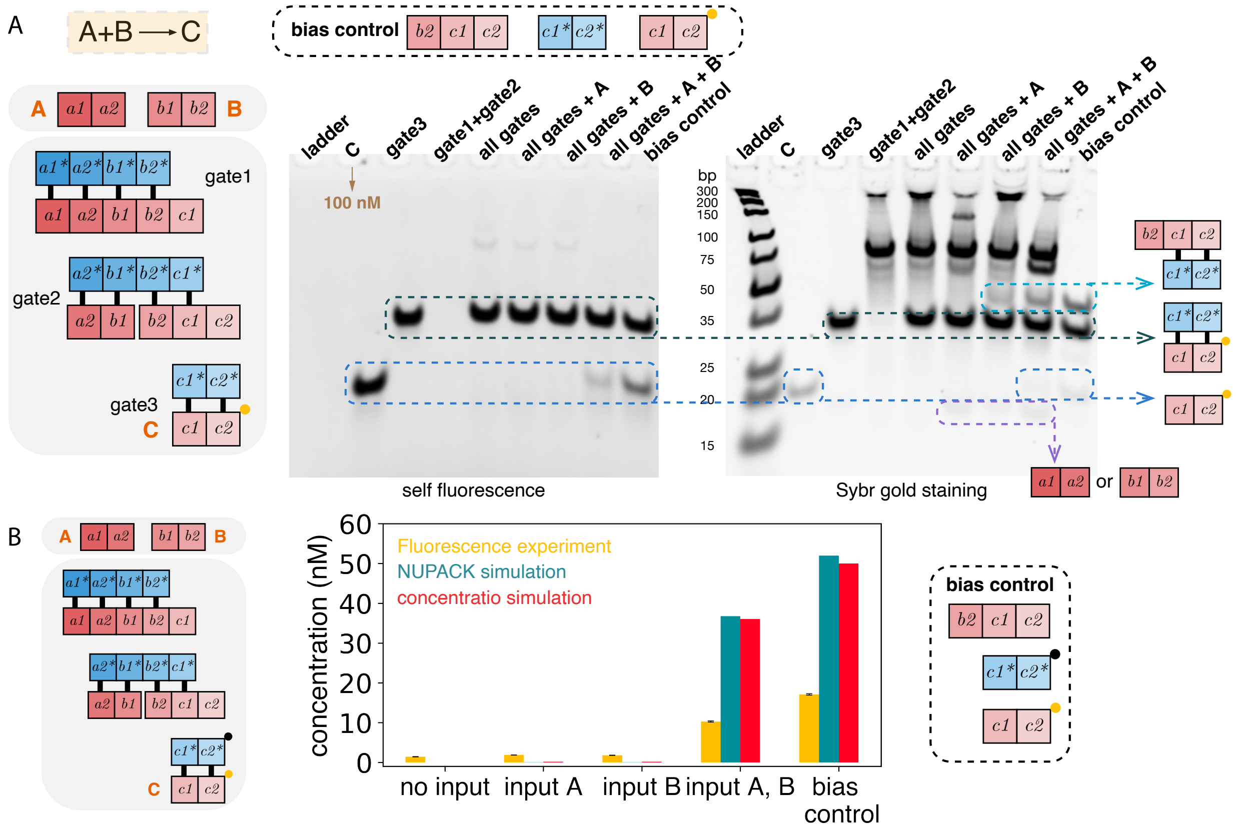


**Figure S3:** The (A) 15% native PAGE gel and (B) fluorescence results for signal propagation circuit $A + B \to C$. The quantitative analysis of the unstained gel is in Figure 2B. The fluorescence experiment introduces an extra quencher (black dot) in the starred molecule in gate3 ($c_1^* c_2^*$). All the molecules have initial concentration 100 nM prior to annealing. The NUPACK simulation has the maximum number of strands in a complex set to be 3, and includes the "leak" complex with 6 strands. The concentratio simulation uses the default parameters. The fluorescence signal is normalized to the signal of 100 nM fluorphore labeled output. The bias control experiment shows that there is preference for $c_1^* c_2^*$ to bind with $C$ compared to $b_2 c_1 c_2\}$. It is not clear what causes the discrepancy, but this could lead to lower than expected output concentration.

#### S3.3 Circuit for $C \to A + B$

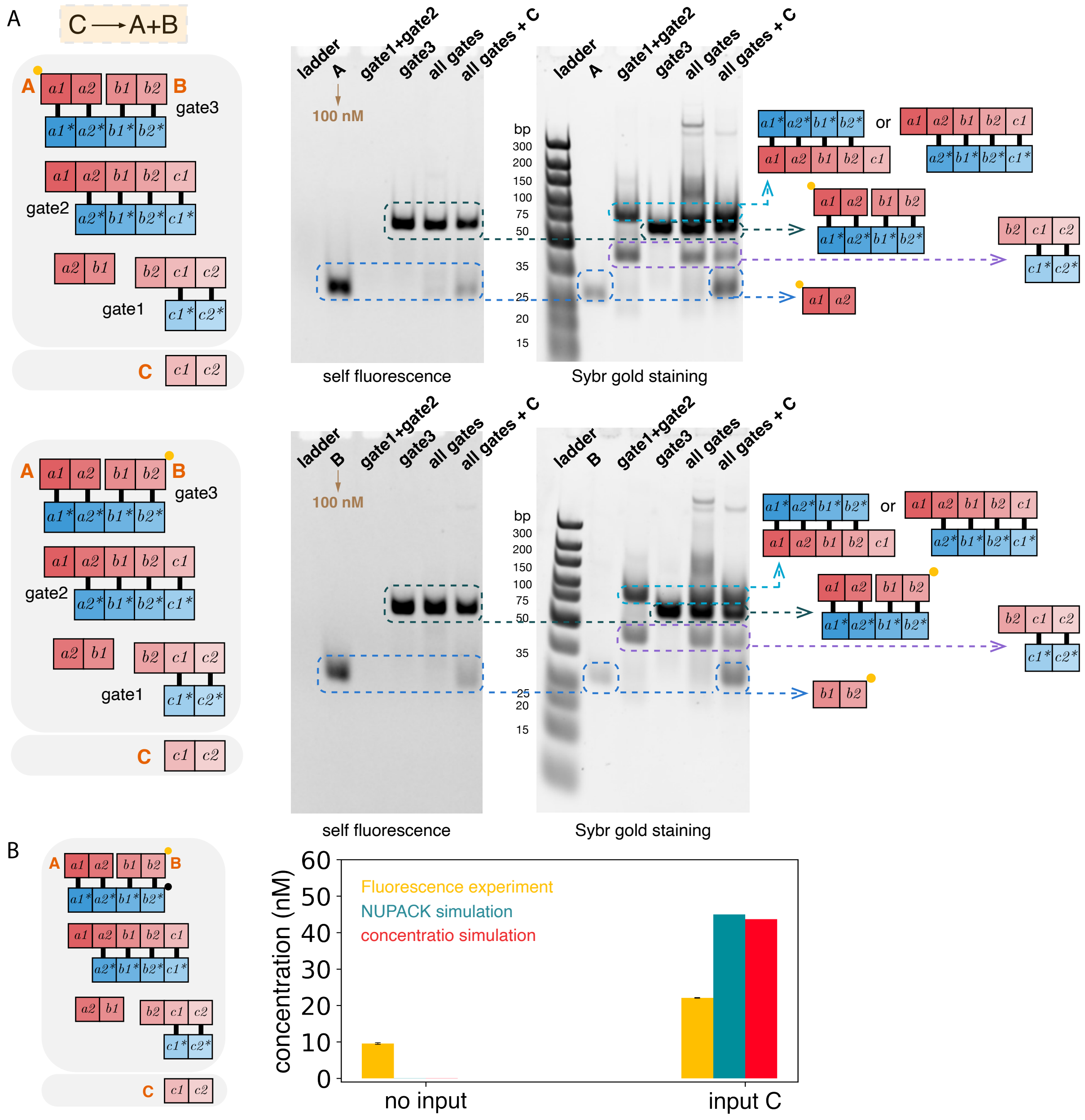


**Figure S4:** The (A) 10% native PAGE gel and (B) fluorescence results for signal propagation circuit $C \to A + B$. The top and bottom pairs of gels have output $A$ and output $B$ labeled by a fluorophore, respectively. The quantitative analysis of the unstained gel is in Figure 2C. The fluorescence experiment introduces an extra quencher (black dot) in the starred molecule in gate3 ($a_1^* a_2^* b_1^* b_2^*$). All the molecules have initial concentration 100 nM prior to annealing. The fluorescence signal is normalized to the signal of 100 nM fluorphore labeled output. The NUPACK simulation has the maximum number of strands in a complex set to be 3, and includes the "leak" complex with 6 strands. The concentratio simulation uses the default parameters.

#### S3.4 Numerical analysis of equilibrium concentrations

The TBN model abstracts thermodynamic favorability through combinatorially characterizing the stable configurations (i.e., configurations maximizing the number of separate complexes among saturated configurations). When every species is at a single copy, we expect the same probability of being in each of the stable configurations, all else being equal. However, in bulk experiments where species have many copies, the extra entropic contribution due to combinatorial entropy can bias the different stable configurations differently, even if their free energies are identical.[2] Combinatorial entropy can also push the system toward configurations that are considered not stable according to the TBN model.

While it remains an open question to better theoretically characterize the equilibrium concentration of bulk TBN systems, numerical simulation tools can be used to analyze even large systems. In this section, we study the expected concentrations of output in the $A + B \rightarrow C$ signal propagation system (alternatively described as an AND gate), and composition of such modules, using numerical analysis with the `concentrat.io` web tool via the available API [9]. Note that this tool operates at the domain-level unlike the more detailed base-level analysis of tools like Nupack, and can thus handle much larger TBN systems such as the composition of multiple signal propagation modules analyzed below.

We first characterize the effect of proportionally scaling all concentrations. In the $A + B \rightarrow C$ signal propagation system, *leak* is defined to be the production of free output $C$ in the absence of at least one input $A$ or $B$. *Correct output* is defined to be the production of free output $C$ when both inputs are present. Leak decreases the number of separate complexes by one (i.e., incurs a cost of a unit of entropy; see Section S3.1). Since the driving force of a unit of entropy increases with decreasing concentration, we expect better fidelity at lower concentration. Note that this effect is expected in both bulk and single molecule systems (where concentration is the inverse of the volume). Numerical analysis shown in Figure S5 confirm that the relative amount of leak decreases with decreasing concentration. All non-input strands are present at equal concentration $c$ nM as shown on the x-axis. The inputs $A$ and $B$ are either also present at concentration $c$ nM (intended output case), or one or both of them are absent (leak cases). In each case, we compute the concentration of free output $C$, and plot the ratio of this concentration in the leak case divided by the intended output case. The ratio (super-linearly) approaches $0$ with decreasing $c$. The ratio of the amount of intended output to $c$ is independent of $c$ (not shown).

Next we focus on the bias toward different stable configurations in the bulk regime, and how the bias can be shifted via the combinatorial entropy. As pointed out in the main text, when both inputs $A$ and $B$ are added, there are three stable configurations, only one of which has free $C$. In Figure S6 we show that, for a fixed concentration of inputs, increasing the concentration of non-input strands leads to more output. Note that increasing the concentration of non-input strands also increases leak, although it remains small even at high concentrations.

Finally we consider systems of multiple $A + B \rightarrow C$ modules together, where the output of the upstream module is input to a downstream module. In Figure S7 we consider a linear topology of the cascade as shown. As the depth of the cascade $k$ (i.e., number of AND gates) increases, the amount of correct output decreases slightly but approaches an asymptote. This is expected via similar considerations as analyzed in [10] (Section "Asymptotic Completion Level of Translator

[2] This is the entropy accounting for different ways to form a complex from constituent indistinguishable parts. For example, complex $A{:}B$ will be present in higher concentration than complex $A{:}C$ if $[B] > [C]$ and both complexes have the same free energy of formation.

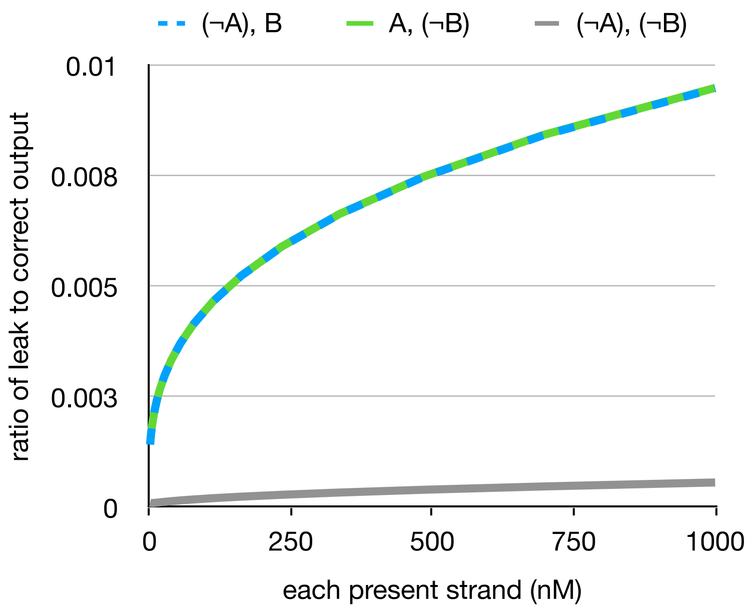


**Figure S5:** Scaling of the ratio of leak to correct output with changing concentration (nM). The y-axis shows the ratio of free output $C$ given one or neither input versus free output $C$ given both inputs. Three types of leak are considered: without input $A$, without input $B$, without both inputs. The same concentration of each present strand is used as shown on the x-axis.

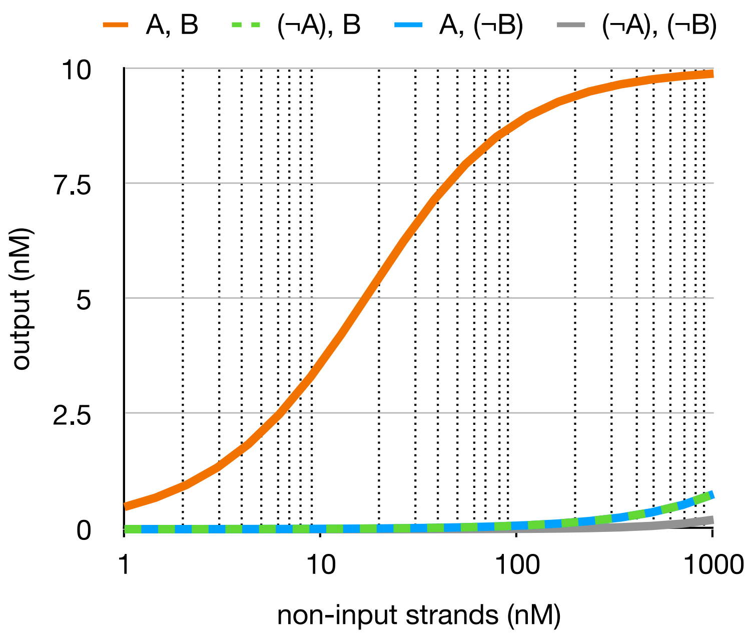


**Figure S6:** Amount of free output $C$ as a function of the concentration of non-input strands. Four cases are considered: correct output (both inputs are present), and three cases of leak. The inputs ($A$, $B$) when present are at 10 nM each.

Cascades"). Nonetheless, the amount of leak increases with $k$, shrinking the gap between correct output and leak. Thus the fidelity of long cascades would require further decreasing concentration (making a unit of entropy stronger) or redesigning the system to increase the number of units of entropy penalty to leak (see Discussion in main text).

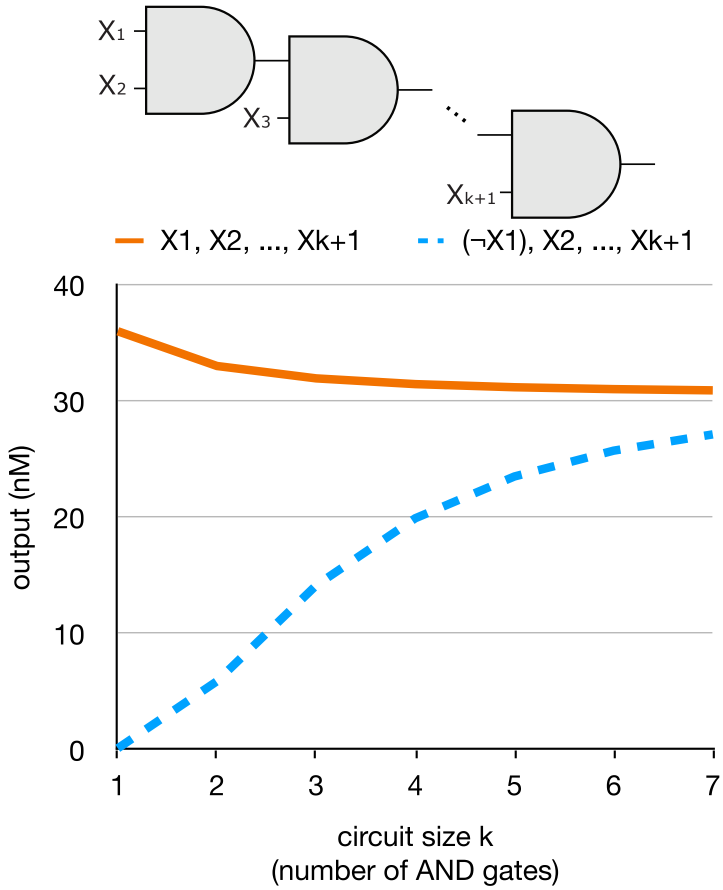


**Figure S7:** Scaling of correct output and leak with increasing cascade length of a linear cascade of $A + B \to C$ modules (shows as AND gates). All concentrations are 100nM. Maximum complex size parameter was chosen to be 6.

#### S3.5 Sequences

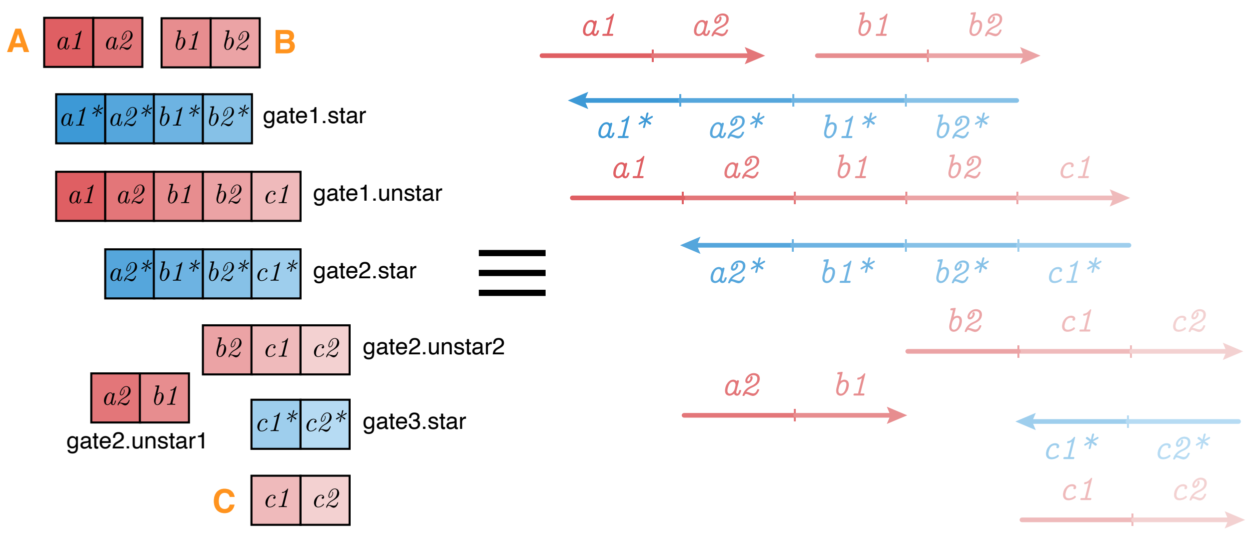


**Figure S8:** The DNA strands for the signal propagation circuit corresponding to the box abstraction.

"gate1.star", "gate1.unstar", and "gate2.star" were ordered PAGE purified. All other unmodified oligos and fluorophore or quencher labeled oligos were ordered HPLC purified.

The sequences are named based on the circuit $A + B \rightarrow C$.

| | |
|---|---|
| A | TTTTCCATTTCCAAC CCAAAACCCTTTTAC |
| B | CATATCCTATCCCAC CATCATCAATACACC |
| C | TCTCACAAACTAACT ACTCTTCTTCTTCTC |
| gate1.star | GGTGTATTGATGATG GTGGGATAGGATATG GTAAAAGGGTTTTGG GTTGGAAATGGAAAA |
| gate1.unstar | TTTTCCATTTCCAAC CCAAAACCCTTTTAC CATATCCTATCCCAC CATCATCAATACACC TCTCACAAACTAACT |
| gate2.star | AGTTAGTTTGTGAGA GGTGTATTGATGATG GTGGGATAGGATATG GTAAAAGGGTTTTGG |
| gate2.unstar.1 | CCAAAACCCTTTTAC CATATCCTATCCCAC |
| gate2.unstar.2 | CATCATCAATACACC TCTCACAAACTAACT ACTCTTCTTCTTCTC |
| gate3.star | GAGAAGAAGAAGAGT AGTTAGTTTGTGAGA |
| A-F | /56-ROXN/TTTTCCATTTCCAAC CCAAAACCCTTTTAC |
| B-F | CATATCCTATCCCAC CATCATCAATACACC/3Rox_N/ |
| C-F | TCTCACAAACTAACT ACTCTTCTTCTTCTC/3Rox_N/ |
| gate1.star-Q | /5IAbRQ/GGTGTATTGATGATG GTGGGATAGGATATG GTAAAAGGGTTTTGG GTTGGAAATGGAAAA |
| gate3.star-Q | /5IAbRQ/GAGAAGAAGAAGAGT AGTTAGTTTGTGAGA |

#### S4 Domain order for unpseudoknotted implementation

As shown in Figure S9, converting the abstract TBN notation to DNA molecules may require organizing domain order in molecules to make it geometrically feasible. Although pseudoknotted structures could be geometrically feasible, here we mainly consider unpseudoknotted implementation because the geometrical structure is more straightforward. We first show that certain TBN configurations cannot be implemented with unpseudoknotted structures. Then we show an algorithm to generate domain order of each molecule for unpseudoknotted implementation.

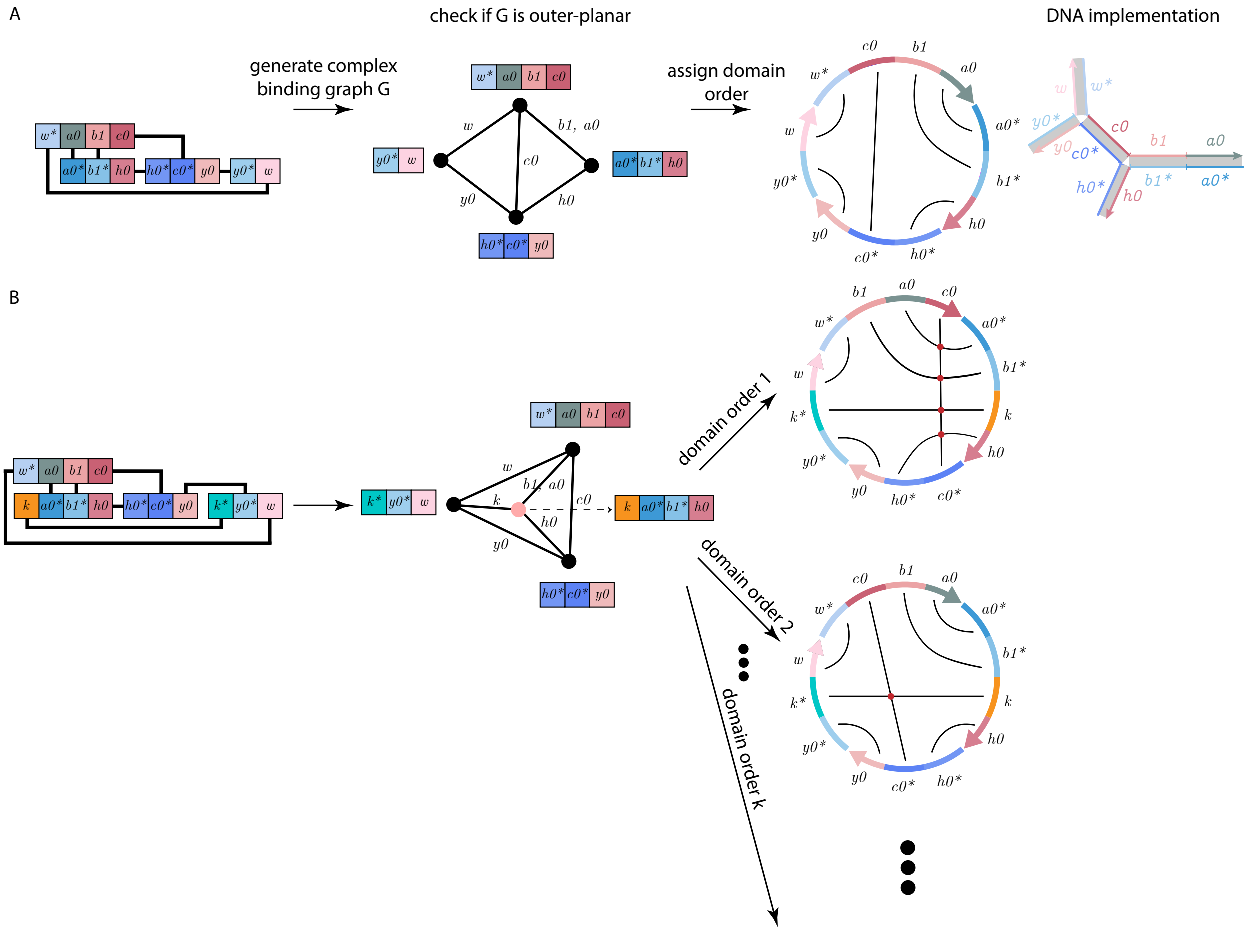


**Figure S9:** Converting the abstract TBN notation to DNA molecules: First a complex binding graph is generated according to the bonds between the monomers. If the complex binding graph is outer-planar, as the example shown in (A), then there exists a unpseudoknotted implementation; If the complex binding graph is not outer-planar (but it could be planar), as the example shown in (B), then there does not exist any unpseudoknotted implementation. Domain orders on each monomers for unpseudoknotted implementation can be decided through ordering the domains counterclockwise on each vertex of the complex binding graph.

We first define the TBN abstraction more formally, to prove that our method for finding unpseudoknotted implementations is general. Given a set of *domains*, e.g., $\{a, b, c\}$, there is an implicit set of complementary-binding *starred domains*, e.g., $\{a^*, b^*, c^*\}$. A monomer is a multiset of domains, e.g., $\{a, a, b^*\}$. As a multiset, we do not consider the domains to be in a particular order. We define a complex as a multiset of monomers. As a multiset, a complex does not explicitly denote which domains within a complex are bound.

For our algorithm to find unpseudoknotted implementations of a complex, we require the user's additional input as to which pairs of monomers are bound by at least one or more domains. Thus the user specifies a *complex binding graph* where each vertex represents a monomer and each edge represents the presence of one or more bonds between two monomers.

Next we define the notion of pseudoknots. Fixing an ordering of domains on every strand, a polymer graph (drawing) [2] can be used to represent secondary structures of a complex. A *polymer graph drawing* of a circular ordering of strands is defined as follows: place the domains of all strands around the circumference of a circle consistent with the order of the strands and the domains within strands. Then connect the bound complementary domains with straight lines. Different circular orderings of the strands represent distinct polymer graph drawings. In this representation, a secondary structure is *pseudoknotted* if every circular strand ordering corresponds to a polymer graph drawing with crossing lines; otherwise, the secondary structure is *unpseudoknotted*. It has been shown that for an unpseudoknotted secondary structure, there is exactly one circular strand ordering corresponding to a polymer graph drawing with no crossing lines [2, Theorem 2.1] (assuming the structure is connected—i.e., the complex binding graph is connected).

Given an ordering of domains on strands and an ordering of strands, it is easy to check whether the corresponding polymer graph drawing has crossing lines. However, there are exponentially many such orderings, making it not obvious how to find such an ordering if it exists. The following claim characterizes the easy-to-check property of the complex binding graph that allows us to order domains to yield unpseudoknotted secondary structures. We say a graph is *outerplanar* if it has a planar embedding (i.e., drawing with no crossing edges) where all vertices are on the outer face.

**Theorem S4.1.** *There exists an unpseudoknotted implementation of a complex if and only if its complex binding graph is outer-planar.*

*Proof.* We prove each direction separately.

($\implies$): If a complex is unpseudoknotted, by reference [2], there exists exactly one circular permutation of a connected polymer graph with no crossing lines. Take this strand order as the order for the vertices in the complex binding graph, and all the vertices are on the outer face. Since there is no crossing lines in the polymer graph for this circular permutation of strand order, there are no crossing edges in the complex binding graph. Thus this complex binding graph is outer-planar.

($\impliedby$): Since all the vertices are on the outer face of the complex binding graph, there exists a circular order of the vertices without crossing edges. Take this circular order of the vertices as the strand order for the complex in the polymer graph. Since each vertex represents a strand, the domain order in the strand can be determined by ordering the bonds in each edge connected to the vertex counterclockwise. Since the complex binding graph is outerplanar and there is no crossing edge, the polymer graph with this specific strand order will not be crossing. □

If no outerplanar embedding exists for the complex binding graph, then the molecular implementation must be pseudoknotted. There exist polynomial time algorithms which check if a graph is outer-planar and, if so, output an outer-planar embedding [11, 12]. Since the domain order in all strands can be determined in polynomial time as well, the algorithm for determining whether a domain order resulting in an unpseudoknotted DNA implementation exists, and giving the domain order if it does, is polynomial time.

If a pseudoknotted structure is unavoidable, introducing poly T could overcome geometric constraints without carefully designing domain length for certain structures to form.

### S5 Seeded-assembling circuits

#### S5.1 The role of cap molecules and the $w$ domain

While it is clear how the gate molecules encode the truth table of their corresponding logic gate in their domains, this section explains the role of the cap molecules and the additional $w$ and $w^*$ domain on the input monomer and the output monomer. Note that the formal proof of correctness is given in Section S5.2.

First, we note that the cap monomers prevent self-assembly of circuits which do not include the input monomer. If the TBN did not have cap monomers, in fig. S10A the configuration would not be saturated, and saturation would require assembling gate monomers into at least a partial circuit without the input monomer. With the cap monomers in the TBN, assembling gate monomers without an input monomer (fig. S10B, highlighted in red) yields a configuration with fewer complexes than leaving them separate (fig. S10A).

The role of the $w$ domain can be described as follows. Beginning with the configuration in fig. S10C, which has all gate monomers alone with their cap monomers, one can assemble gates onto the input monomer while displacing their cap monomers without changing the number of separate complexes. Due to the $w$ domain, which forces an output monomer to bind to the input monomer to satisfy saturation, the addition of the last gate in this process displaces its own cap along with the output monomer's cap, yielding an extra unit of entropy, as shown by the final configuration in fig. S10A.

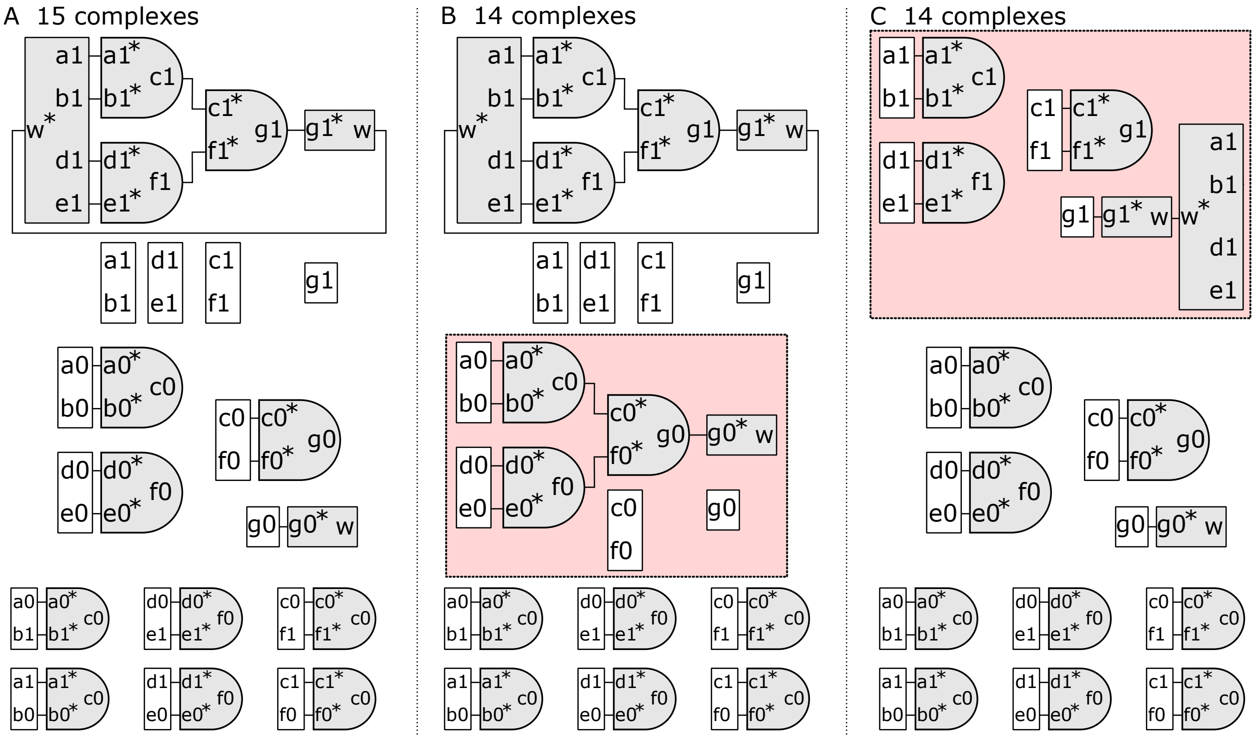


**Figure S10:** A seeded-assembling circuit TBN consisting of three AND gates. The input, gate, and output monomers are colored gray, and the cap monomers are colored white. (A) A stable configuration of the TBN. (B) Assembling a circuit without an input monomer (red) yields fewer complexes than leaving them separate as in part A. (C) Disassembling the desired circuit while maintaining saturation (red) yields fewer complexes than leaving the circuit assembled as in part A.

##### S5.2 Proof that stable configurations have correct circuits

In this section we provide an abridged version of the proof shown in [13], that any stable configuration of a seeded-assembling circuit construction includes a full and correct circuit assembly.

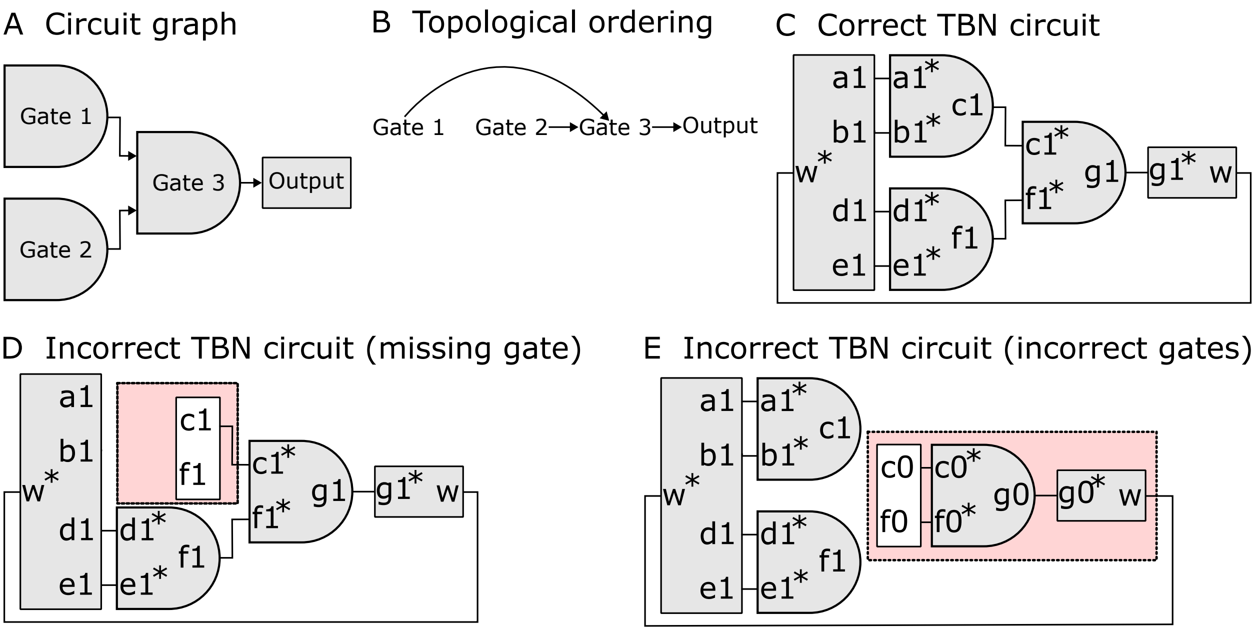


**Figure S11:** Helpful illustrations for understanding the proof that stable configurations have correct circuits. (A) A feedforward circuit represented as a directed acyclic graph. (B) A topological ordering of the directed acyclic graph from part A. (C) The notion of "correct" circuit complex for this TBN system (see Figure S10 a for the full TBN). (D) A complex which is incorrect because a gate is missing. **(E)** A complex which is incorrect because the gates used do not have correct logic. In both cases **(D)** and **(E)**, the proof argues that a cap (white) must be in the circuit to maintain saturation.

First note that a feedforward circuit can be represented by a directed acyclic graph (see Figure S11A), and the graph can be topologically sorted to give an ordering on gates where gate $j$ takes inputs from gates $i < j$ and gives outputs to gates $k > j$ (see Figure S11B). Also note that starred domains are limiting: for any domain $x$, the system has at least as much of $x$ as $x^*$. So any $x^*$ must be bound in any saturated configuration, and thus in any stable configuration. By construction, a gate's starred domains can only bind to an upstream gate, an input, or a cap. For the proof, we call the output molecule a gate (it is last in the topological ordering). Figure S11C shows a correct circuit for comparison to incorrect ones. Figure S10A may also help to understand the proof as a reference of a full configuration, which is the same TBN as the examples in Figure S11.

In all of the following claims, we assume that the configurations referenced are saturated (though not necessarily stable).

**Claim S5.1.** *If a gate is missing or incorrect in the complex with the input, the configuration has some cap and the input in the same complex.*

*Proof.* Consider the complex with the input. If there is a missing gate, let $i$ be the latest missing gate with respect to the topological ordering. See Figure S11D for an example configuration with a missing gate. If gate $i$ is the output molecule, the $w^*$ domain on the input is not bound, contradicting saturation. Otherwise, a gate $k > i$ with a starred domain designed to bind to the missing gate $i$ must be bound to a cap. If a gate is incorrect, let $i$ be the earliest incorrect gate with respect to the topological ordering. See Figure S11E for an example configuration with an incorrect gate. Since gates $j < i$ are correct, at least one of gate $i$'s starred domains (the incorrect domain) cannot bind to a gate. It also cannot bind to the input, so it must be bound to a cap. In either case, there is a cap in the same complex as the input molecule. □

**Claim S5.2.** *Any gate molecule must be in a complex with either a cap or the input molecule.*

*Proof.* Let $i$ be the index of the gate in question. For $i$ such that the gates bind to the input, note that the starred domains can only bind to the input or a cap. We will induct on $i$. Consider a gate $i$ such that gates $j < i$ satisfy the claim. The gate $i$'s starred domains can only bind to a cap or one of the gates $j < i$, or the input. If bound to some gate $j$, because that gate satisfies the claim, the gate $i$ also satisfies the claim. □

This immediately implies:

**Claim S5.3.** *The number of separate complexes is at most the number of complexes with a cap or input molecule.*

Finally the following claim establishes the correctness of the construction:

**Theorem S5.4.** *Any stable configuration has a complete, correct circuit.*

*Proof.* By Claim S5.1, if a gate is missing or incorrect, the configuration has some cap and the input in the same complex. Thus by Claim S5.3, the number of separate complexes is at most the number of caps. By the existence of a configuration with a number of complexes equal to the number of caps plus one (the desired configuration, see Figure S10A), the missing or incorrect configuration is not stable. □

##### S5.3 Data Normalization

Experiment data were normalized according to the signals of two control samples of signal level "1" and "0":

- Normalized signal level "1" (Negative control): strands "Out0" and "Out1" mixed together, which represents the signal levels where no fluorescence is quenched.
- Normalized signal level "0" (Positive control): strand "Out0" or "Out1" mixed with one strand with quencher, which represents the signal levels where each of the fluorophore is quenched.

##### S5.4 Kinetic pathway and input flipping

We show that the NAND gate responds to input triggers within an hour. The logic gate can be correctly reset if the stable configuration has the input bound with its complement. After resetting, the NAND gate can be triggered with a different input.

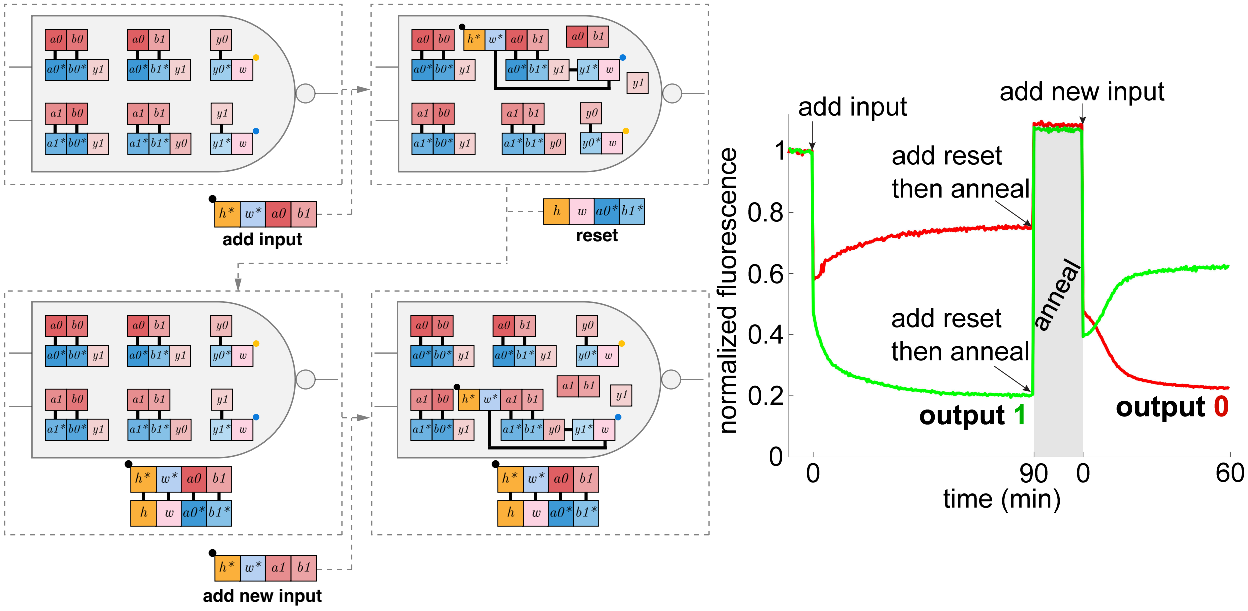


**Figure S12:** Kinetics of a NAND gate in response to input 01, and annealed for resetting, and kinetics of flipping to input 11. 30% ethanol was mixed in the reaction buffer to increase the kinetic rate.

#### S5.5 Deletion

The correctness of the circuit is designed by having the configuration with correct computation with one more separate complexes than all other configurations with incorrect computation, assuming all the configurations have the same enthalpic energy (the same number of bonds). However, the assembled structure formed from correct computation loses stacking energy at the junctions. This lose of enthalpic energy could lead to correct computation being unfavorable. Figure S13A shows an example with input 010. Assuming all other molecules are the same, the configuration with the correct circuit assembly is not energetically favored, and the difference of free energy is 8-12 kcal/mol.

In order to balance the loss of stacking energy at junctions in the correct circuit assembly, we decreased the enthalpic energy in the left configuration by deleting some nucleotides in the cap strands. NUPACK predicts that deleting one nucleotides increases the free energy by around 4 kcal/mol, thus we tried deleting one or two nucleotides in the cap strand. Experimental results show that having deletion improves the circuit correctness and we then chose 2 deletions to test all 8 inputs.

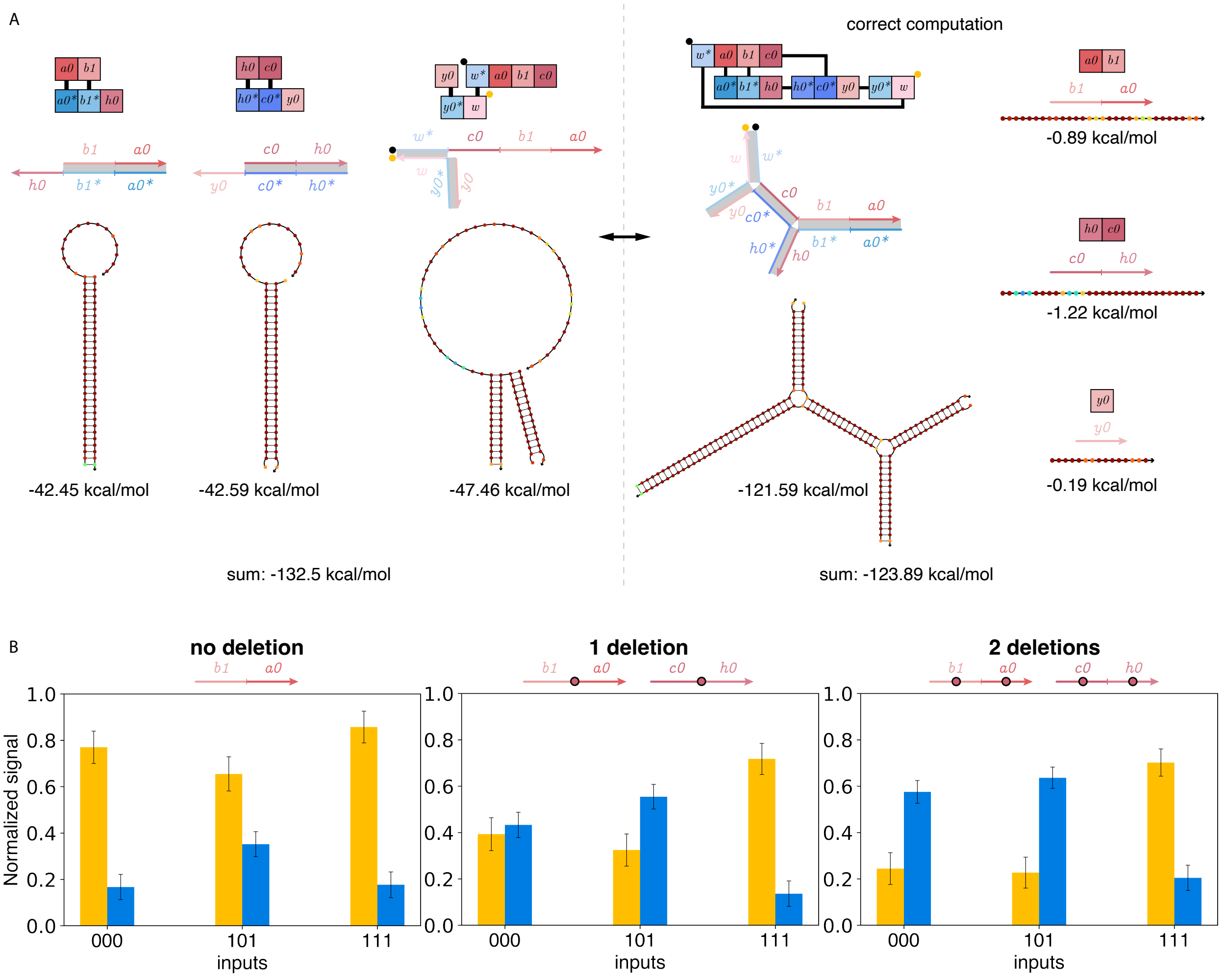


**Figure S13:** Deletion of a few nucleotides on cap strands improves computation correctness. (A) Box abstraction and DNA strand of two different configurations in the TBN model with the same number of bonds but different number of separate complexes. The NUPACK minimum free energy structure and complex free energy of the complexes are also shown. (B) Experiments on the 2-layer AND circuits with different numbers of deletion on some cap strands.

#### S5.6 Sequences

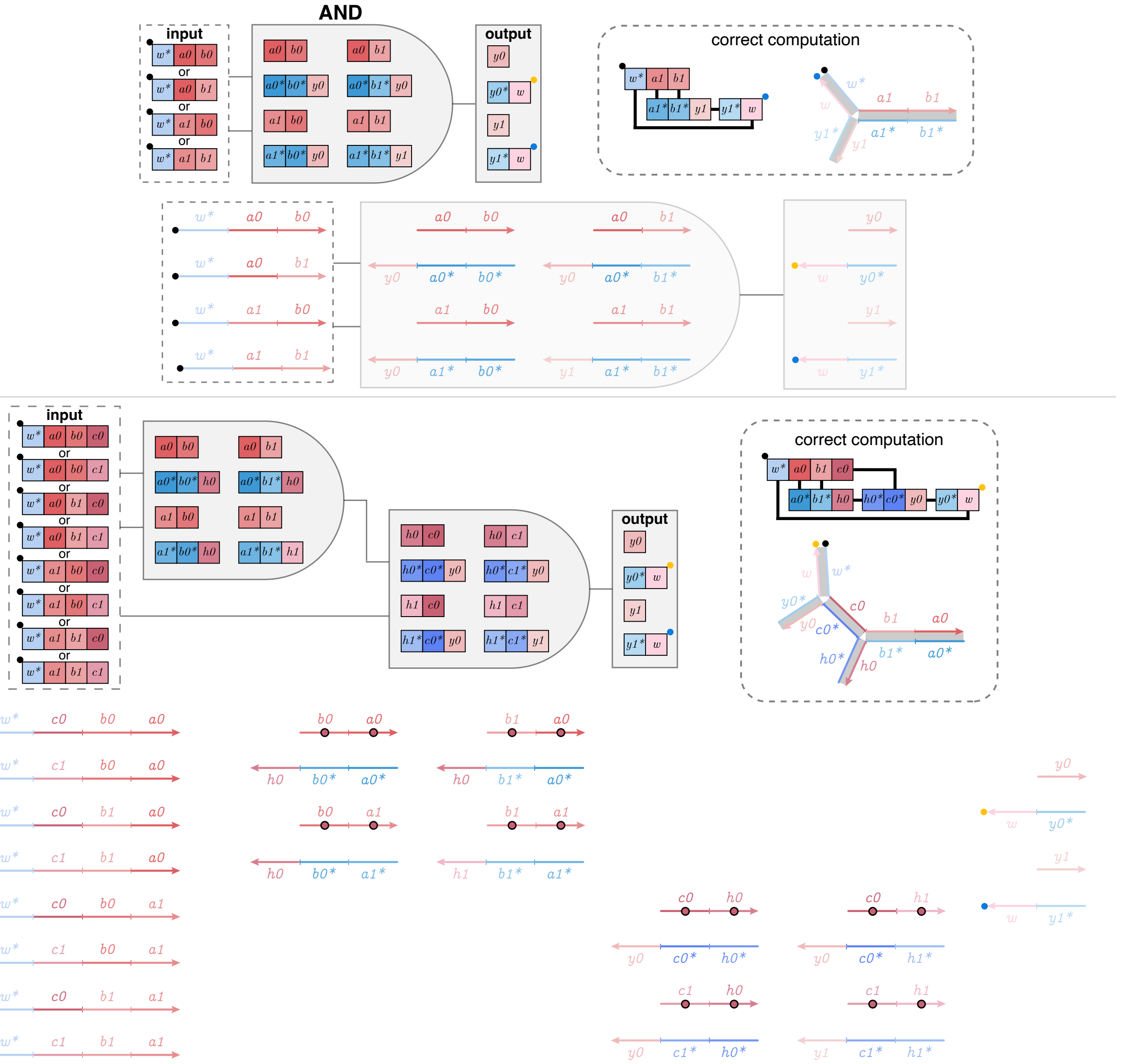

**Figure S14:** The DNA strands for the seeded-assembling circuits corresponding to the box abstraction. The red circle with black outline indicates a deletion in the DNA strand.

The single AND and NAND gates share the same input, output and cap sequences. All the oligos were ordered HPLC purified.

**The single gate**

| | |
|---|---|
| Input00 | /5IAbRQ/AAGGTTTGGTTGATT ATGGGAAGAGATATG GTGTTAAGAAGGGAA |
| Input01 | /5IAbRQ/AAGGTTTGGTTGATT ATGGGAAGAGATATG TAGTAATTGGGATGG |
| Input10 | /5IAbRQ/AAGGTTTGGTTGATT ATTGGTAGGAGTTAT GTGTTAAGAAGGGAA |
| Input11 | /5IAbRQ/AAGGTTTGGTTGATT ATTGGTAGGAGTTAT TAGTAATTGGGATGG |
| AND00 | TTCCCTTCTTAACAC CATATCTCTTCCCAT TGGATAGATGGGTAA |
| AND01 | CCATCCCAATTACTA CATATCTCTTCCCAT TGGATAGATGGGTAA |
| AND10 | TTCCCTTCTTAACAC ATAACTCCTACCAAT TGGATAGATGGGTAA |
| AND11 | CCATCCCAATTACTA ATAACTCCTACCAAT GTAGTGTGATGAATA |
| NAND00 | TTCCCTTCTTAACAC CATATCTCTTCCCAT GTAGTGTGATGAATA |
| NAND01 | CCATCCCAATTACTA CATATCTCTTCCCAT GTAGTGTGATGAATA |
| NAND10 | TTCCCTTCTTAACAC ATAACTCCTACCAAT GTAGTGTGATGAATA |
| NAND11 | CCATCCCAATTACTA ATAACTCCTACCAAT TGGATAGATGGGTAA |
| Out0 | TTACCCATCTATCCA AATCAACCAAACCTT/3ATTO550N/ |
| Out1 | TATTCATCACACTAC AATCAACCAAACCTT/3ATTO647NN/ |
| Gate00Cap | ATGGGAAGAGATATG GTGTTAAGAAGGGAA |
| Gate01Cap | ATGGGAAGAGATATG TAGTAATTGGGATGG |
| Gate10Cap | ATTGGTAGGAGTTAT GTGTTAAGAAGGGAA |
| Gate11Cap | ATTGGTAGGAGTTAT TAGTAATTGGGATGG |
| Out0Cap | TGGATAGATGGGTAA |
| Out1Cap | GTAGTGTGATGAATA |

**The 2-layer AND cascade (The nucleotides in red were deleted when ordered)**

| | |
|---|---|
| Input000 | /5IAbRQ/GGTGGTTGATAATTG ACTTCCCAAATCTTT CAACATCTACACCTT AATCCACTCACTATC |
| Input001 | /5IAbRQ/GGTGGTTGATAATTG TCAAATCCCTTTTCT CAACATCTACACCTT AATCCACTCACTATC |
| Input010 | /5IAbRQ/GGTGGTTGATAATTG ACTTCCCAAATCTTT TCACAAACAAAACAA AATCCACTCACTATC |
| Input011 | /5IAbRQ/GGTGGTTGATAATTG TCAAATCCCTTTTCT TCACAAACAAAACAA AATCCACTCACTATC |
| Input100 | /5IAbRQ/GGTGGTTGATAATTG ACTTCCCAAATCTTT CAACATCTACACCTT ACACATAACCTCATT |
| Input101 | /5IAbRQ/GGTGGTTGATAATTG TCAAATCCCTTTTCT CAACATCTACACCTT ACACATAACCTCATT |
| Input110 | /5IAbRQ/GGTGGTTGATAATTG ACTTCCCAAATCTTT TCACAAACAAAACAA ACACATAACCTCATT |
| Input111 | /5IAbRQ/GGTGGTTGATAATTG TCAAATCCCTTTTCT TCACAAACAAAACAA ACACATAACCTCATT |
| AND1-00 | GATAGTGAGTGGATT AAGGTGTAGATGTTG TCTCCTCTCTATTCA |
| AND1-01 | GATAGTGAGTGGATT TTGTTTTGTTTGTGA TCTCCTCTCTATTCA |
| AND1-10 | AATGAGGTTATGTGT AAGGTGTAGATGTTG TCTCCTCTCTATTCA |
| AND1-11 | AATGAGGTTATGTGT TTGTTTTGTTTGTGA ACTAACCATTTACCC |
| AND2-00 | TGAATAGAGAGGAGA AAAGATTTGGGAAGT CTTTCATCCTCCATA |
| AND2-01 | TGAATAGAGAGGAGA AGAAAAGGGATTTGA CTTTCATCCTCCATA |
| AND2-10 | GGGTAAATGGTTAGT AAAGATTTGGGAAGT CTTTCATCCTCCATA |
| AND2-11 | GGGTAAATGGTTAGT AGAAAAGGGATTTGA ACTAACTTTCCAACC |
| Out0 | TATGGAGGATGAAAGCAATTATCAACCACC/3ATTO647NN/ |
| Out1 | GGTTGGAAAGTTAGTCAATTATCAACCACC/3ATTO550N/ |
| Out0Cap | CTTTCATCCTCCATA |
| Out1Cap | ACTAACTTTCCAACC |
| AND1-00Cap | CAACATCTACACCTT AATCCACTCACTATC |
| AND1-01Cap | TCACAAACAAAACAA AATCCACTCACTATC |
| AND1-10Cap | CAACATCTACACCTT ACACATAACCTCATT |
| AND1-11Cap | TCACAAACAAAACAA ACACATAACCTCATT |
| AND2-00Cap | ACTTCCCAAATCTTT TCTCCTCTCTATTCA |
| AND2-01Cap | TCAAATCCCTTTTCT TCTCCTCTCTATTCA |
| AND2-10Cap | ACTTCCCAAATCTTT ACTAACCATTTACCC |
| AND2-11Cap | TCAAATCCCTTTTCT ACTAACCATTTACCC |
| AND1-00Cap-d1 | CAACATCTACACCTT AATCCACTCACTATC |
| AND1-01Cap-d1 | TCACAAACAAAACAA AATCCACTCACTATC |
| AND1-10Cap-d1 | CAACATCTACACCTT ACACATAACCTCATT |
| AND1-11Cap-d1 | TCACAAACAAAACAA ACACATAACCTCATT |
| AND2-00Cap-d1 | ACTTCCCAAATCTTT TCTCCTCTCTATTCA |
| AND2-01Cap-d1 | TCAAATCCCTTTTCT TCTCCTCTCTATTCA |
| AND2-10Cap-d1 | ACTTCCCAAATCTTT ACTAACCATTTACCC |
| AND2-11Cap-d1 | TCAAATCCCTTTTCT ACTAACCATTTACCC |
| AND1-00Cap-d2 | CAACATCTACACCTT AATCCACTCACTATC |
| AND1-01Cap-d2 | TCACAAACAAAACAA AATCCACTCACTATC |
| AND1-10Cap-d2 | CAACATCTACACCTT ACACATAACCTCATT |
| AND1-11Cap-d2 | TCACAAACAAAACAA ACACATAACCTCATT |
| AND2-00Cap-d2 | ACTTCCCAAATCTTT TCTCCTCTCTATTCA |
| AND2-01Cap-d2 | TCAAATCCCTTTTCT TCTCCTCTCTATTCA |
| AND2-10Cap-d2 | ACTTCCCAAATCTTT ACTAACCATTTACCC |
| AND2-11Cap-d2 | TCAAATCCCTTTTCT ACTAACCATTTACCC |

### S6 Size-controllable concatemers

**Claim S6.1.** *When the ratio of the molecular counts of the molecules is $n : m$, the only complex in the stable configuration contains the least common multiple of $m$ and $n$ ($\mathrm{LCM}(m, n)$) of domains $x^*$.*

*Proof.* By having the ratio of the molecular counts $n : m$, the total number of domain $x$ is equal to that of domain $x^*$, thus any saturated configuration of the system does not have any complex with unbound domains. Let $L$ be the complex with $\mathrm{LCM}(m, n)$ number of domain $x$ and $x^*$. $L$ is the smallest complex with no unbound domains. Because all bonds of $L$ are bound, any saturated complex larger than $L$ could separate $L$ from itself and gain one unit of entropy. Thus the stable configuration which contains the maximum number of complexes only has the bounded size complexes $L$. □

#### S6.1 Side product

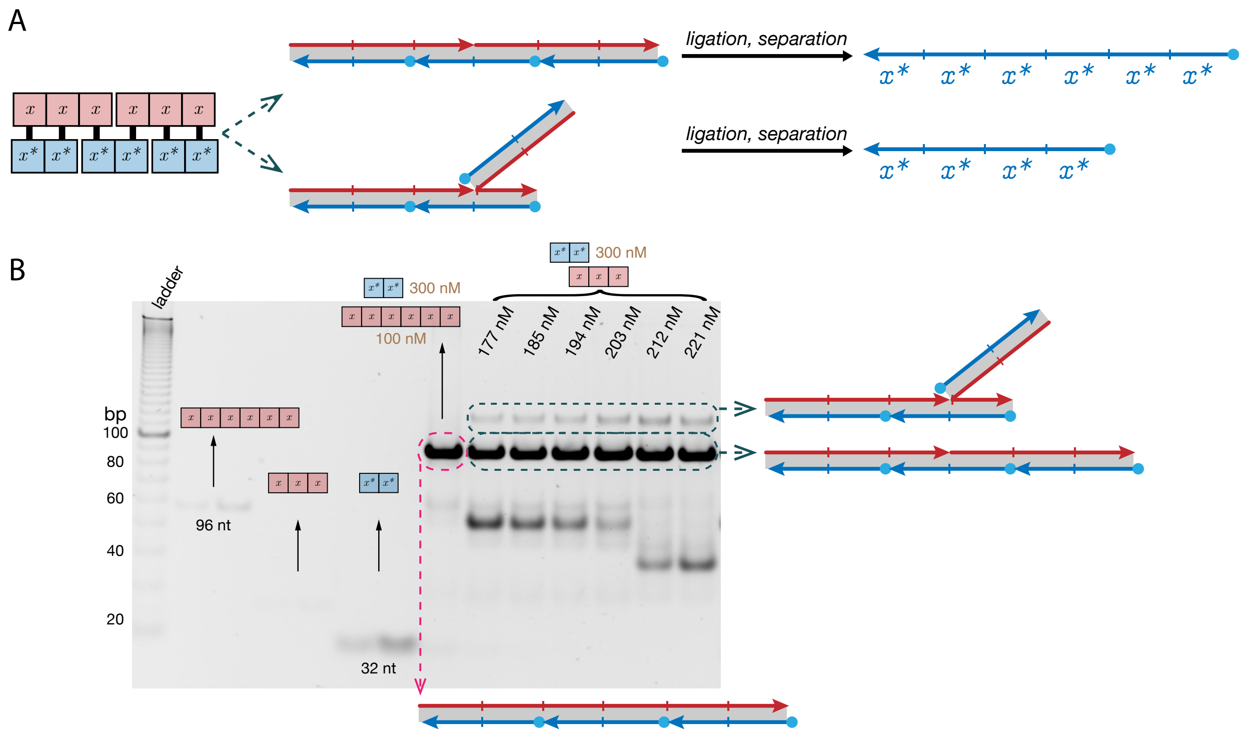


**Figure S15:** Side product for the number pair $(3, 2)$. (A) A possible reaction pathway for generating a side product. Annealing can lead to a branched complex instead of a linear complex. (B) Full gel with controls showing in Figure 5B. The pink lane is a control showing the gel location of the linear product.

##### S6.2 Concatemers with substrates of different sizes

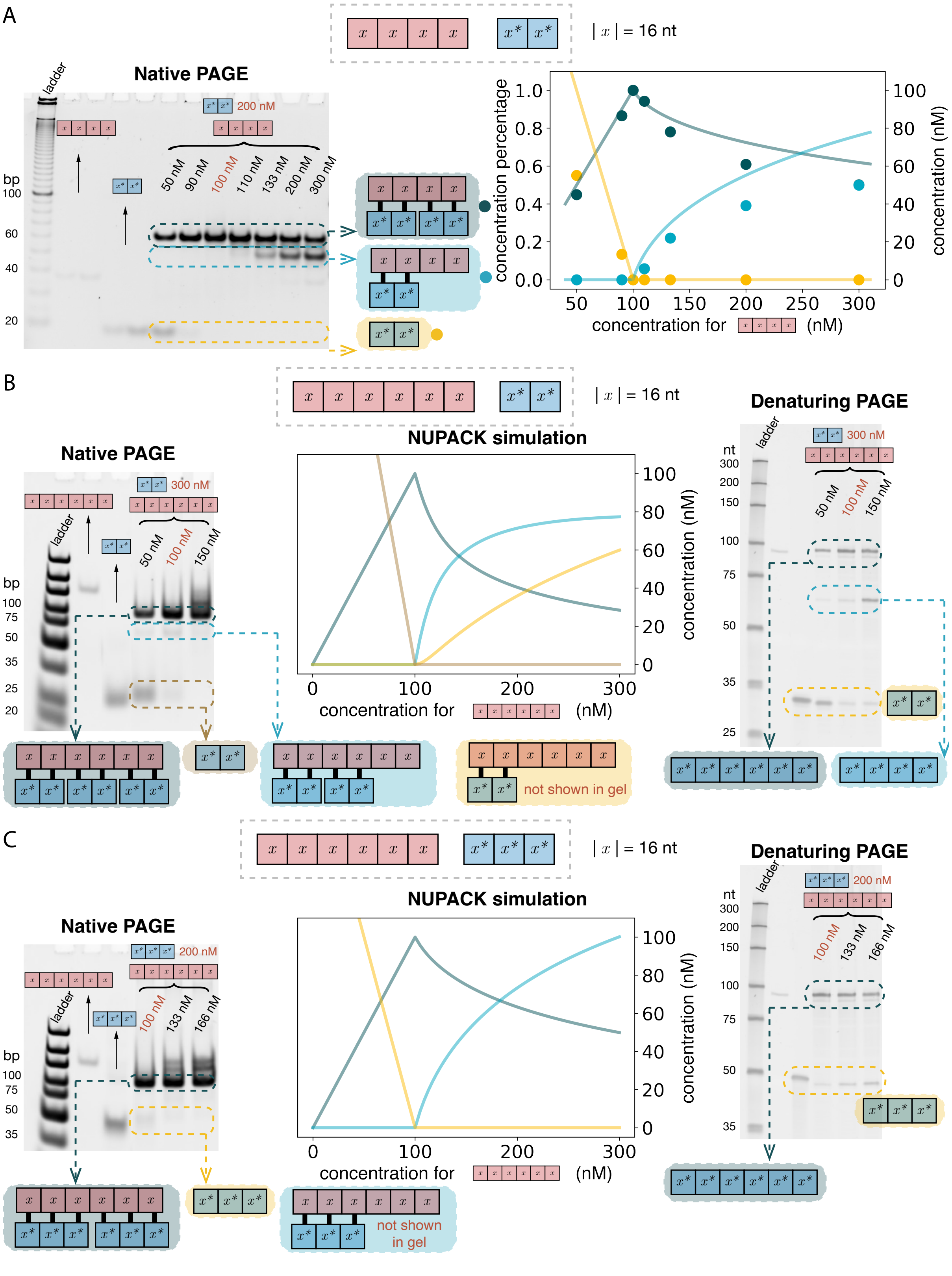


**Figure S16:** Size-controllable concatemers with the size of the substrates not mutually prime (A) $xxxx$ and $x^*x^*$, (B) $xxxxxx$ and $x^*x^*$ and (C) $xxxxxx$ and $x^*x^*x^*$.

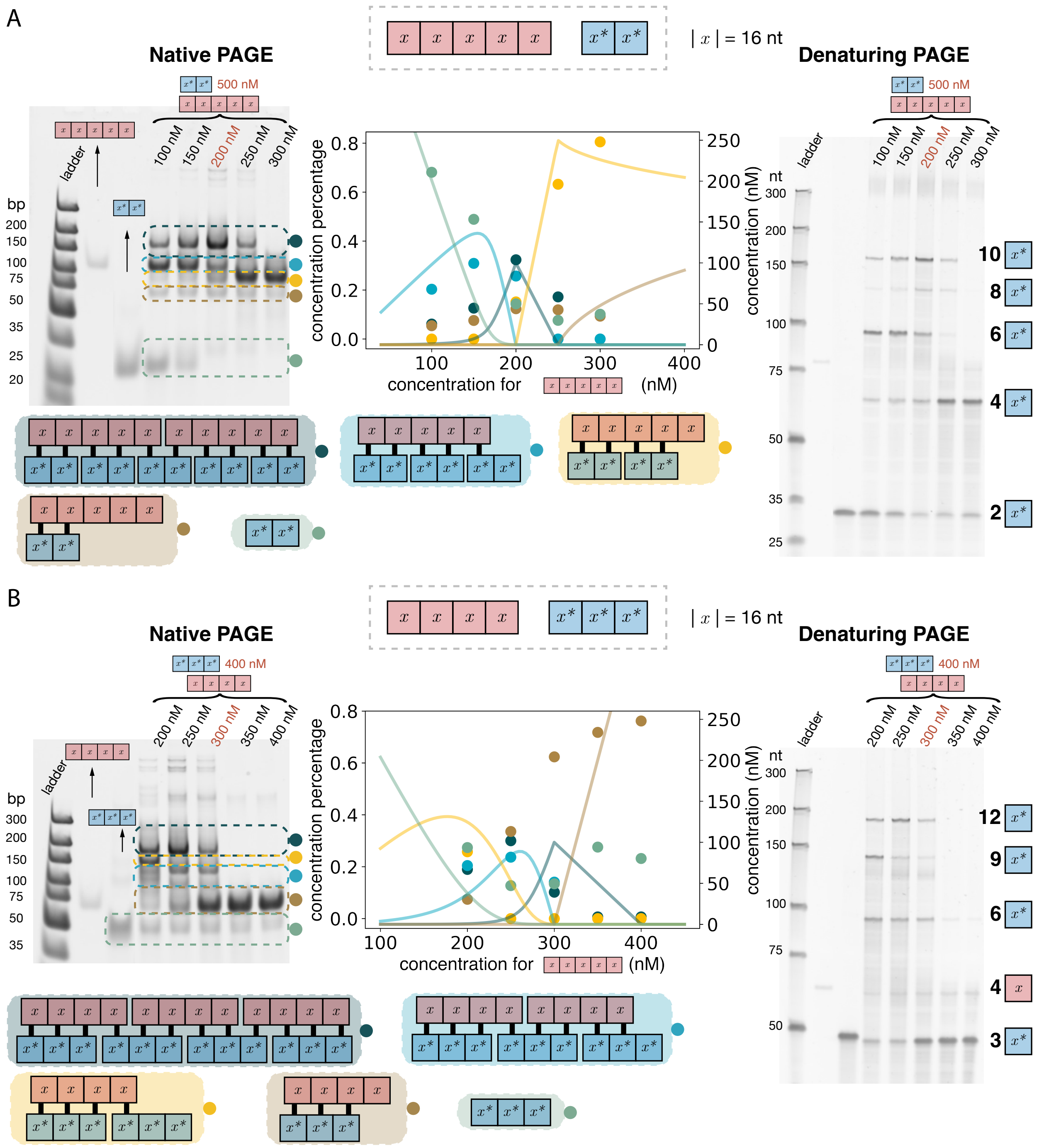


**Figure S17:** Size-controllable concatemers with the size of the substrates mutually prime (A) $xxxxx$ and $x^*x^*$ and (B) $xxxx$ and $x^*x^*x^*$.

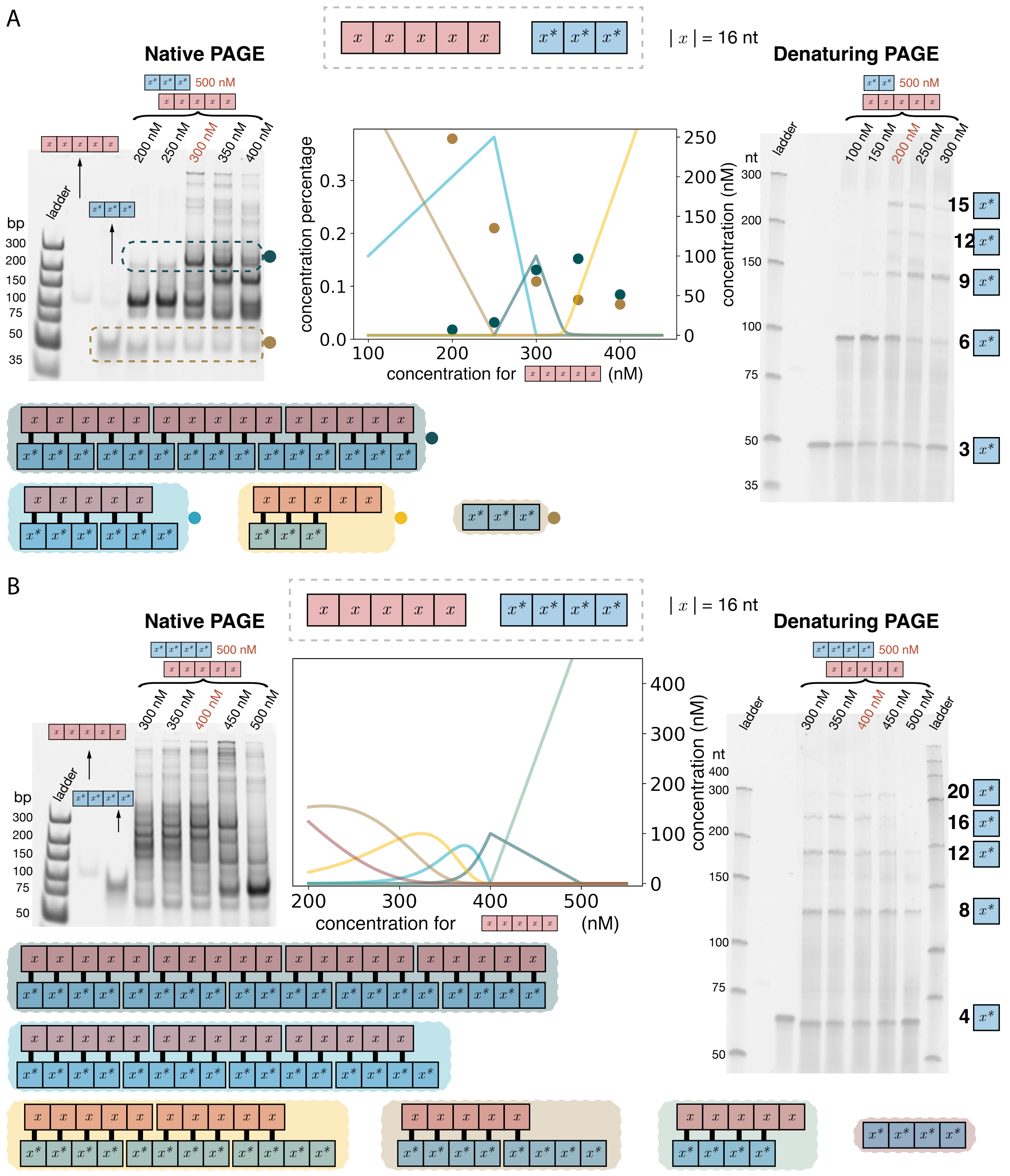


**Figure S18:** Size-controllable concatemers with the size of the substrates mutually prime (A) $xxxxx$ and $x^*x^*x^*$ and (B) $xxxxx$ and $x^*x^*x^*x^*$.

##### S6.3 Sequences

$xxx$ and $xxxx$ were ordered PAGE purified. $x^*x^*$, $x^*x^*x^*$ and $x^*x^*x^*x^*$ have phosphate labeled and were ordered HPLC purified. $xxxxx$ and $xxxxxx$ were ordered as PAGE Ultramer.

| | |
|---|---|
| $xxx$ | CTCTCTCTTAACCCAC CTCTCTCTTAACCCAC CTCTCTCTTAACCCAC |
| $xxxx$ | CTCTCTCTTAACCCAC CTCTCTCTTAACCCAC CTCTCTCTTAACCCAC CTCTCTCTTAACCCAC |
| $xxxxx$ | CTCTCTCTTAACCCAC CTCTCTCTTAACCCAC CTCTCTCTTAACCCAC CTCTCTCTTAACCCAC CTCTCTCTTAACCCAC |
| $xxxxxx$ | CTCTCTCTTAACCCAC CTCTCTCTTAACCCAC CTCTCTCTTAACCCAC CTCTCTCTTAACCCAC CTCTCTCTTAACCCAC CTCTCTCTTAACCCAC |
| $x^*x^*$ | /5Phos/GTGGGTTAAGAGAGAG GTGGGTTAAGAGAGAG |
| $x^*x^*x^*$ | /5Phos/GTGGGTTAAGAGAGAG GTGGGTTAAGAGAGAG GTGGGTTAAGAGAGAG |
| $x^*x^*x^*x^*$ | /5Phos/GTGGGTTAAGAGAGAG GTGGGTTAAGAGAGAG GTGGGTTAAGAGAGAG GTGGGTTAAGAGAGAG |
